# Aberrant expression of collagen type X in solid tumor stroma is associated with EMT, immunosuppressive and pro-metastatic pathways, bone marrow stromal cell signatures, and poor survival prognosis

**DOI:** 10.1101/2024.11.13.621984

**Authors:** Elliot H.H. Famili-Youth, Aryana Famili-Youth, Dongfang Yang, Ayesha Siddique, Elizabeth Y. Wu, Wenguang Liu, Murray B. Resnick, Qian Chen, Alexander S. Brodsky

## Abstract

**Background:** Collagen type X (ColXα1, encoded by *COL10A1*) is expressed specifically in the cartilage-to-bone transition, in bone marrow cells, and in osteoarthritic (OA) cartilage. We have previously shown that ColXα1 is expressed in breast tumor stroma, correlates with tumor-infiltrating lymphocytes, and predicts poor adjuvant therapy outcomes in ER^+^/HER2^+^ breast cancer. However, the underlying molecular mechanisms for these effects are unknown. In this study, we performed bioinformatic analysis of *COL10A1*-associated gene modules in breast and pancreatic cancer as well as in cells from bone marrow and OA cartilage. These findings provide important insights into the mechanisms of transcriptional and extracellular matrix changes which impact the local stromal microenvironment and tumor progression.

**Methods:** Immunohistochemistry was performed to examine collagen type X expression in solid tumors. WGCNA was used to generate *COL10A1*-associated gene networks in breast and pancreatic tumor cohorts using RNA-Seq data from The Cancer Genome Atlas. Computational analysis was employed to assess the impact of these gene networks on development and progression of cancer and OA. Data processing and statistical analysis was performed using R and various publicly-available computational tools.

**Results:** Expression of *COL10A1* and its associated gene networks highlights inflammatory and immunosuppressive microenvironments, which identify aggressive breast and pancreatic tumors and contribute to metastatic potential in a sex-dependent manner. Both cancer types are enriched in stroma, and *COL10A1* implicates bone marrow-derived fibroblasts as drivers of the epithelial-to-mesenchymal transition (EMT) in these tumors. Heightened expression of *COL10A1* and its associated gene networks is correlated with poorer patient outcomes in both breast and pancreatic cancer. Common transcriptional changes and chondrogenic activity are shared between cancer and OA cartilage, suggesting that similar microenvironmental alterations may underlie both diseases.

**Conclusions:** *COL10A1*-associated gene networks may hold substantial value as regulators and biomarkers of aggressive tumor phenotypes with implications for therapy development and clinical outcomes. Identification of tumors which exhibit high expression of *COL10A1* and its associated genes may reveal the presence of bone marrow-derived stromal microenvironments with heightened EMT capacity and metastatic potential. Our analysis may enable more effective risk assessment and more precise treatment of patients with breast and pancreatic cancer.

**Research Highlights:**

- ColX highlights features of EMT in breast and pancreatic cancer
- ColX gene modules are immunosuppressive and pro-metastatic
- ColX-associated gene networks contribute to sex differences in pancreatic cancer
- ColX-positive fibroblasts define more aggressive tumors with poorer survival
- ColX is emerging as a biomarker for bone marrow-derived cells in cancer

## BACKGROUND

The tumor microenvironment profoundly influences cancer progression and aggression through direct and indirect interactions between neoplastic cells and surrounding structures such as the extracellular matrix (ECM), whose remodeling plays a critical role in tumor growth, invasion, and metastasis^1^. The ECM encompasses a broad array of glycoproteins, collagens, and proteoglycans, as well as affiliated regulators and secreted factors; together, these exhibit a multitude of structural and signaling functions across diverse biological contexts and their alteration contributes to the development of pathological states ranging from fibrotic diseases to cancer^2,3^. Collagens are the most abundant protein component of the ECM and the composition of both major and minor collagens has been shown to vary substantially across different cancer types; additionally, numerous collagens have been identified as biomarkers associated with molecular alterations and overall survival in cancers of diverse primary tissues^4^. The composition and distribution of collagens within the local ECM is largely driven by fibroblasts, which influence the processes of inflammation and angiogenesis through regulation of the ECM^5^, although other cells may also play a role in the production and degradation of collagens in disease states. Fibroblastic activity appears to be an important driver of disease across diverse tumor types, but the full composition and function of the ECM in cancer remains uncertain.

Collagen type X (*COL10A1*, ColX) is a non-fibrillar collagen synthesized specifically by hypertrophic chondrocytes to regulate matrix mineralization, stiffness, and metabolism^6^. ColX promotes the cartilage-to-bone transition in skeletal development, is highly expressed by bone marrow stromal cells (BMSCs), and becomes progressively elevated in articular cartilage during the development of osteoarthritis (OA)^6–9^. OA pathogenesis has been associated with senescence of mesenchymal stromal cells, chondrocyte death, calcification and degradation of the extracellular matrix, and angiogenic invasion^10–12^. Previously we found that ColX is expressed in breast tumor tissue, the first time that ColX was shown to be highly expressed in non-skeletal tissues^13–15^. Prior studies by our group have shown that ColX is not only expressed in many types of breast tumors, but is also associated with overall survival outcomes for ER^+^/HER2^+^ breast tumors in particular^13,15^. However, the mechanism by which ColX is involved in tumor progression and treatment outcomes remains unknown. Pancreatic ductal adenocarcinoma (PDAC) remains a highly lethal malignancy due to its aggressive nature and the paucity of effective treatment options; such tumors are notable for their high fractions of desmoplastic stroma which contributes significantly to drug and immune resistance^16^. Complex interactions between PDAC cancer cells and surrounding stromal features such as activated fibroblasts and collagens play a major role in aggressive, treatment-refractory disease^16,17^. Thus, one approach to improve our understanding of stromal impacts in cancer is to identify key ECM features which drive the development, survival, and progression of such tumors.

Core environmental factors which influence tumor outcomes include stromal composition, blood vessel density, and infiltrating immune cells^18^. The complicated interplay between resident and foreign host cells, the extracellular matrix, and molecular signals all contribute to primary tumor treatment responses. Recent studies have suggested that the stromal fractions of breast and pancreatic tumors feature significant proportions of cancer-associated fibroblasts (CAFs), which exhibit substantial heterogeneity, originate from both the local biome and differentiated bone marrow-derived mesenchymal stromal cells, and contribute significantly to patient prognosis and response to therapy^19,20^. Given ColX’s important role in cartilage development and the bone marrow niche^21^, along with its dysregulated expression across both OA cartilage and solid tumors, we sought to characterize its pathophysiologic role in cancer through bioinformatic analysis of *COL10A1*-expressing cancer and non-cancer cells. We hypothesized that similar stromal microenvironments across certain cancers, bone marrow, and OA cartilage are defined by ColX and its associated gene networks, which may contribute to molecular mechanisms underlying tumor progression. In this study, we defined gene co-expression modules to characterize pathways and microenvironmental components related to and correlated with ColX expression in breast and pancreatic tumors from The Cancer Genome Atlas (TCGA). By analyzing ColX expression in BMSCs and OA cartilage cells, we found notable common biological pathways in both cancer and BMSCs, thereby linking these two types of stromal cells. Characterization of ColX expression networks and their pathological mechanisms will improve understanding of aggressive disease states and offer opportunities for devising future therapies.

## METHODS

### Immunohistochemistry and ColXα1 expression scoring

Two PAAD samples were tested compared to breast tumor observations. One stage 3 and one stage 4 sample were evaluated for ColXα1 protein expression as follows. Four-micron sections were cut from formalin-fixed paraffin-embedded tissue blocks, heated at 60°C for 30 minutes, deparaffinized, rehydrated, and subjected to antigen retrieval by heating the slides in epitope retrieval buffer in a water bath at 95°C for 45 minutes. The slides were then incubated with either mouse monoclonal antibodies or rabbit polyclonal antibodies for 30 minutes at room temperature in a DAKO Autostainer. Anti-ColXα1 (1:100, eBioscience/Affymetrix, Clone X53) was used for immunohistochemistry (IHC). Immunoreactivity was detected using the DAKO EnVision method according to the manufacturer’s recommended protocol.

### Data analysis and visualization

All data processing and analysis was performed in R (version 4.0.2)^22^ unless otherwise stated. Visualizations were generated in R using the ggplot2 (version 3.3.6)^23^, gplots (version 3.1.3)^24^, and eulerr (version 7.7.0)^25,26^ packages. See Figure 1C for overview of tumor sample datasets and computational tools employed in this study.

**Figure 1:**
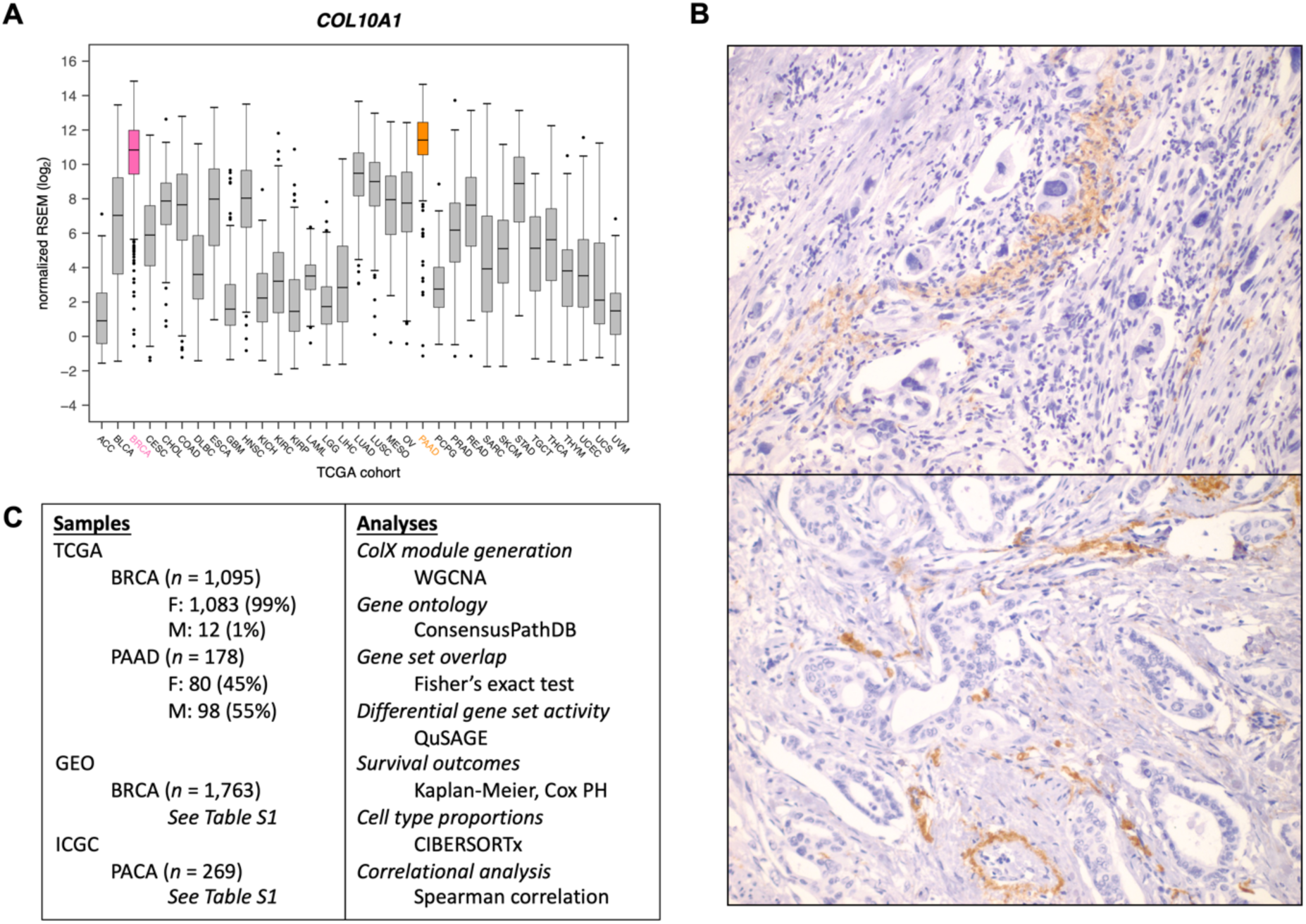
*COL10A1* is highly expressed in breast and pancreatic tumors. **(A)** Expression of *COL10A1* across all TCGA sample cohorts. **(B)** Representative 400x ColXα1 immunohistochemistry staining in pancreatic tumors. **(C)** Outline of study samples and analyses. See Methods and Results sections for details.

### Data acquisition and pre-processing

#### TCGA gene expression data

Log-normalized expression for *COL10A1* across all TCGA cancer types was downloaded from the Broad Institute’s Genome Data Analysis Center FireBrowse portal. Batch-corrected, normalized RNA-Seq-derived RSEM values for breast invasive carcinoma (BRCA, *n* = 1,095 samples) and pancreatic adenocarcinoma (PAAD, *n* = 178 samples) TCGA cohorts were downloaded from the NIH National Cancer Institute Genomic Data Commons^27,28^. Genes with RSEM values < 1 in ≥ 50% of samples and mean RSEM values < 50 overall were defined as “low-expression” and excluded from downstream analysis.

#### Microarray gene expression data

To assess whether RNA-Seq-derived ColX-related modules would be robust to different methodologies of gene expression quantification, microarray datasets with similar sample sizes were selected for comparison to results from the BRCA and PAAD cohorts from TCGA. For breast cancer, raw fluorescence intensity data from a previously-described collection of 12 studies on primary early-stage breast cancer in females was downloaded from the NCBI Gene Expression Omnibus^29^ (GEO) database (*n* = 1,763 samples in total), all of which were expression-profiled on the GPL570 (Affymetrix Human Genome U133 Plus 2.0 Array) platform (see Table S1A for comparison to TCGA data and Table S1B for GSE accessions and number of samples analyzed from each study)^30–42^. For pancreatic cancer, array-based gene expression data for the Australian pancreatic cancer cohort (PACA-AU, *n* = 269 samples) was downloaded from the International Cancer Genome Consortium (ICGC) Data Portal^43^. Pre-processing of microarray data was carried out using the following packages in R: oligo (version 1.52.1)^44^, hgu133plus2.db (version 3.2.3)^45^, AnnotationDbi (version 1.50.3)^46^, tidyverse (version 1.3.0)^47^, WGCNA (version 1.69)^48,49^, and sva (version 3.36.0)^50^. Probe intensity values across each cancer were log-transformed and normalized using the Robust Multichip Average (RMA) quantile method. Probes were mapped to gene IDs based on the GPL570 annotation database, and unmapped or multi-mapping probes were removed. Expression values for multiple probes mapping to the same gene were consolidated using the collapseRows function. Gene expression values for the breast cancer samples were then combined across GEO studies and batch-corrected using the ComBat function. Finally, “low-intensity” expression thresholds were established for each cancer dataset, and all genes with expression values below these thresholds in > 80% of samples were defined as “low-expression” and excluded from downstream analysis.

#### OA gene expression data

Raw RNA-Seq-derived gene counts for 4 knee joint-derived OA cell types (normal cartilage stromal cells/NCSCs, OA mesenchymal stromal cells/OA-MSCs, OA chondrocytes/OACs, bone marrow stromal cells/BMSCs; *n* = 3 each) were sourced from GEO accession GSE176199^10^. Genes exhibiting fewer than 5 counts in ≥ 90% of samples were defined as “low-expression” and excluded from downstream analysis. DESeq2 (version 1.34.0)^51^ was used to normalize raw gene counts and perform differential expression analysis.

For each OA cell type, cell type-specific genes were defined based on the definition of “tissue-enriched’” employed by the Human Protein Atlas (at least four-fold higher mRNA level in a given tissue compared to any other tissues)^52^; i.e., all genes with log_2_-fold change ≥ +2 and adjusted p-value < 0.05 relative to each other cell type.

### Characterization of ColX consensus modules

#### Generation of ColX modules

Correlated gene network modules were generated for each TCGA dataset based on normalized, filtered RSEM values using WGCNA (version 1.69)^48,49^. Briefly, for each dataset, the soft thresholding power *β* was selected to ensure approximately scale-free topology of the gene co-expression network, and signed modules were generated using the blockwiseModules function with parameters deepSplit = 2 and minModuleSize = 30. ColX modules were defined for each dataset separately, comprising all genes which co-modularized with *COL10A1*.

#### Enrichment analysis

Enrichment analysis of ColX modules was performed using the ConsensusPathDB web tool (release 35)^53^. For each cancer, the *over-representation analysis* tool was run to identify all Reactome pathways and gene ontology (GO) terms enriched in the respective ColX module, using as background all genes which were retained after pre-processing the dataset from which the module was generated. Significantly-enriched pathways/terms were identified by FDR-corrected p-value < 0.05.

#### Gene set overlap analysis

Matrisome, hallmark pathway, and Gene Transcription Regulation Database (GTRD) transcription factor target (TFT) gene sets were downloaded from MSigDB^2,54,55^. Overlaps with breast and pancreatic cancer ColX modules were calculated using Fisher’s exact test. Adjusted p-values (p.adj) were computed using the Benjamini-Hochberg (BH) method^56^. Significantly-enriched gene sets were identified by p.adj < 0.05 (p.adj < 0.10 for candidate discovery of enriched TFTs) and odds ratio > 1. Protein-protein interactions of specific transcription factors (TFs) of interest were queried using the STRING database^57^.

#### Module preservation analysis

Preservation of TCGA BRCA and PAAD RNA-Seq-derived ColX modules in corresponding cancer microarray datasets of comparable sample size was assessed using the modulePreservation function following standard WGCNA methodology^58^. The *Z_summary_* statistic, defined as the mean of summarized density preservation statistics and connectivity preservation statistics, was computed for each TCGA-generated module in order to assess the relative preservation of the ColX module. A *Z_summary_* value > 10 was considered to indicate significant module preservation.

### Differential pathway analysis

#### Gene set analysis

Hallmark pathway and GTRD TFT gene sets were downloaded from MSigDB as described above. Immunome gene sets were obtained from The Cancer Immunome Atlas^59^. Cancer-associated fibroblast gene sets were extracted from previously-published datasets and defined as all genes exhibiting ≥ 2-fold increased expression (with p.adj < 0.05) in each fibroblast phenotype of interest relative to all others^19^.

#### Definition of the G.A.M.E. metric

To compare ColX module expression across all tumor samples within each dataset, a ranking metric was defined based on the sample-wise percentage of ColX module genes expressed above their respective median values across all samples (percentage of **G**enes **A**bove-**M**edian **E**xpression, “%G.A.M.E.”). This effectively transformed the unimodal ColX module eigengene (ME) distribution into a bimodal G.A.M.E. distribution with a fixed range between 0 and 1, facilitating clustering of samples into discrete groups (Figure S3). The getJenksBreaks function from the BAMMtools package (version 2.1.10)^60^ was used to divide samples into “low”, “medium”, and “high” G.A.M.E. groups, which were used to proxy ColX module expression for comparisons between subgroups.

#### QuSAGE analysis

Differential activity of gene sets between high and low G.A.M.E. tumor samples were assessed using qusage (version 2.22.0)^61–63^. Log-fold change was used to quantify association with differential ColX module expression (i.e., enrichment). Activation and inhibition of individual gene sets/pathways were defined as positive and negative enrichment relative to the background gene set, respectively. Significance associated with ColX module expression was determined by BH-adjusted p-value < 0.05 and absolute magnitude of enrichment ≥ 25% of the magnitude of activity of the ColX module (both measured relative to the background gene set) within that dataset.

### Survival analysis

#### Cox proportional hazards model analysis

Survival analysis was carried out in R using the survival (version 3.3-1)^64,65^ and survminer (version 0.4.9)^66^ packages. Multivariate Cox proportional hazards models were constructed to assess the statistical dependence of overall survival (OS) and disease-free interval (DFI) on patient age, gender, tumor stage (binarized as stage 1-2 vs. stage 3-4), and ColX tumor signal (assessed as either normalized gene expression, ColX module eigengene (ME), or %G.A.M.E.) for breast and pancreatic cancer cohorts as well as gender-specific pancreatic cancer subcohorts. The proportional hazards assumption was validated for each signal variable by a test of the scaled Schoenfeld residuals. Significant *β* coefficients were determined by a p-value < 0.05, and the per-unit contribution of each significant variable to the overall hazard risk was computed as *e^β^*.

#### Kaplan-Meier survival curve analysis

Differential survival outcomes were assessed using Kaplan-Meier analysis. For each comparison, samples were divided into two groups: a “low” %G.A.M.E. group (defined as above), and a “high” %G.A.M.E. group (all other samples).

#### Tumoral osteoarthritic cell type proportion analysis

CIBERSORTx was utilized to quantify OA cell type proportions for all TCGA tumor samples^67^. The *Create Signature Matrix* tool was used to generate a dimensionally-reduced expression signature (1,978 genes) to differentiate the 4 OA cell types profiled (NCSC, OA-MSC, OAC, BMSC). The *Impute Cell Fractions* tool was subsequently employed to infer the approximate proportions of each OA cell type present in each tumor sample, quantified on an absolute scale. Absolute proportions were scaled by sample-wise total OA cell type proportions to obtain sample-wise relative proportions.

## RESULTS

### ColX-associated gene modules are conserved across different cancers

*COL10A1* mRNA expression was found to be the highest in breast invasive carcinoma (BRCA) and pancreatic adenocarcinoma (PAAD) tumors in TCGA (Figure 1A). We evaluated expression of ColX at the protein level by IHC to determine relative levels of expression. Strong expression was observed in BRCA, as we have previously reported^13–15^. Similar to the patterns observed in breast tumors, ColXα1 IHC revealed a range of mild to strong expression of ColX protein only in the stromal regions of pancreatic tumors (Figure 1B). Consistent with much lower expression of *COL10A1* mRNA, no protein expression was observed in colon adenocarcinoma or stomach adenocarcinoma by IHC. In accordance with these observations, we focused on BRCA and PAAD tumors to evaluate the impact of *COL10A1* on cancer pathophysiology.

To define the role of ColX in breast and pancreatic tumors, we analyzed multiple datasets by a variety of statistical methods (Figure 1C). To identify possible roles and gene networks associated with *COL10A1* in breast and pancreatic tumors, we used weighted gene co-expression network analysis (WGCNA) to generate cancer-specific ColX gene modules in the TCGA datasets for each cancer type. We defined “ColX-associated” genes as all those which were co-modularized with ColX in each dataset, signifying genes whose expression was broadly associated with that of ColX across the patient cohorts. This process yielded ColX modules of 423 and 404 genes across the breast and pancreatic cancer datasets, respectively, with 168 (approximately 40%) of these genes co-modularizing with ColX in both cancers (Figure 2A, Table S2A; see Table S2B for full list of WGCNA modules for each dataset).

**Figure 2:**
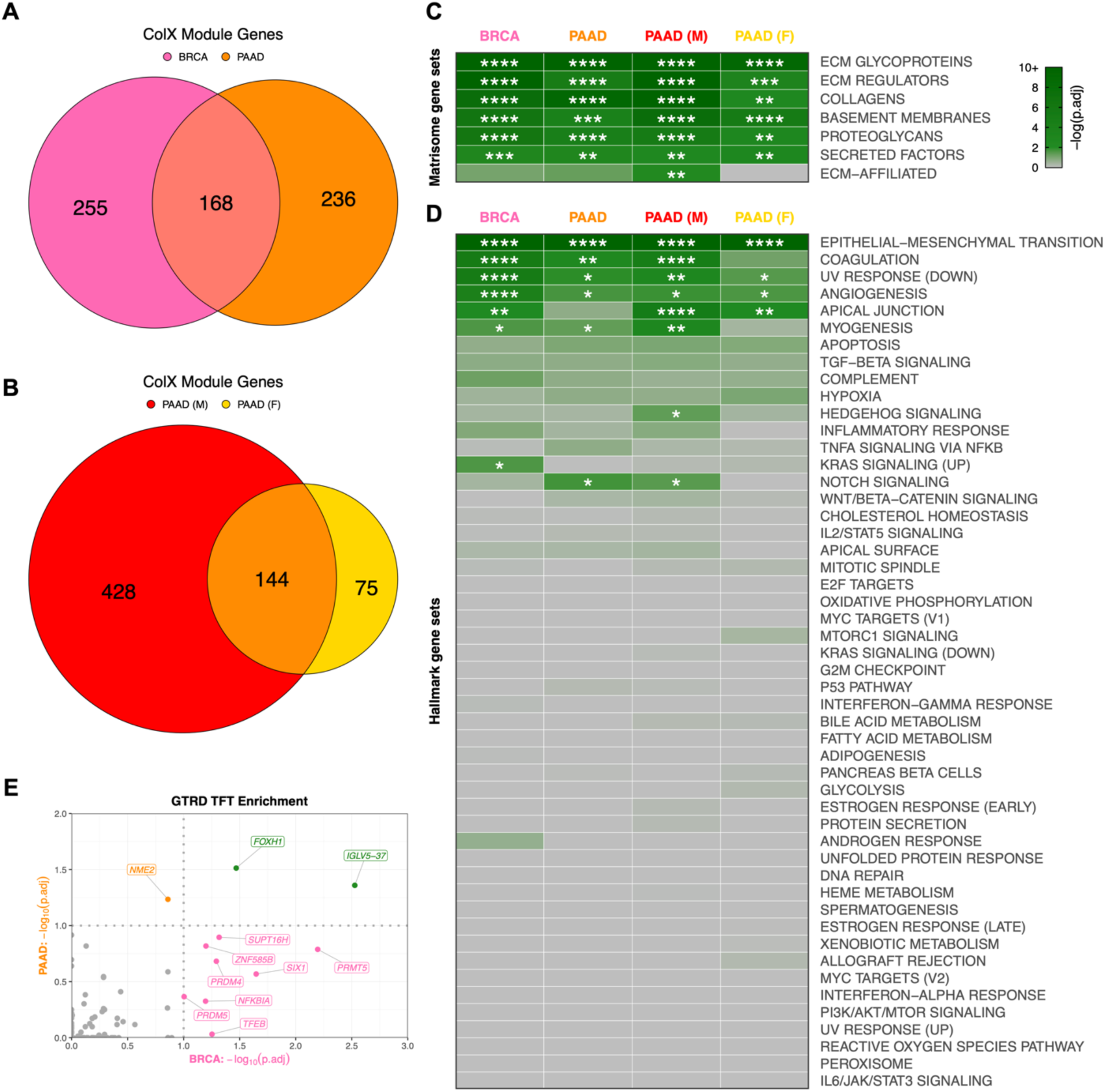
ColX modules are enriched for ECM genes, pro-metastatic pathways, and developmental/regulatory transcription factor targets. **(A and B)** Overlap of genes within ColX WGCNA modules from TCGA **(A)** breast and pancreatic cancer and **(B)** gender-segregated pancreatic cancer datasets. See Table S2A for lists of ColX module genes, Table S2B for full list of WGCNA modules for each dataset, and Table S3 for gene ontology and Reactome pathway enrichment analysis of cancer-specific and overlapping ColX-associated genes. **(C and D)** Enrichment of **(C)** human *in silico* matrisome gene sets^2^ and **(D)** MSigDB hallmark pathway gene sets^54^ within ColX modules. Gene sets within each block are ordered by mean significance rank (by Fisher’s exact test) across all 4 modules. See Figure S2A–D and Table S4 for hallmark pathway enrichment analysis of all WGCNA-inferred modules for each dataset. Significance values: *, p.adj < 0.05; **, p.adj < 0.01; ***, p.adj < 0.001; ****, p.adj < 0.0001. **(E)** Enrichment of Gene Transcription Regulation Database (GTRD) transcription factor (TF) targets^55^ within breast and pancreatic cancer ColX modules. Of note, the TFT gene set attributed to *IGLV5-37* in the GTRD database actually represents targets of the fusion oncoprotein *SS18-SSX*, as described in the text. Dotted lines correspond to p.adj = 0.10. Green labels/points indicate TFs whose targets are enriched in both breast and pancreatic cancer ColX modules. See Table S5A for reported TF functions and lists of overlapping TFTs. Significance values: *, p.adj < 0.05; **, p.adj < 0.01; ***, p.adj < 0.001; ****, p.adj < 0.0001.

To verify the reproducibility of these ColX modules, we performed module preservation analysis on a comparably-sized microarray dataset for each cancer type: a collection of 1,763 primary early-stage breast cancer samples sourced from GEO as well as 269 pancreatic cancer samples from the PACA-AU cohort. Both TCGA RNA-Seq-derived ColX modules were highly conserved in their corresponding microarray cancer datasets; the BRCA ColX module was the #2 most highly preserved among all 31 BRCA gene modules and the PAAD ColX module was the #1 most highly preserved among all 42 PAAD gene modules. This signifies the relevance of these cancer-specific gene sets across diverse patient cohorts (Figure S1).

### ColX-associated gene networks are enriched for ECM and developmental ontologies

To probe the biological importance of these ColX modules in the context of breast and pancreatic cancer, we performed gene ontology (GO) and Reactome pathway enrichment analysis to identify functions of ColX modules in each dataset (Tables S3, S4). As anticipated, GO terms relating to extracellular matrix and collagen function were highly enriched across both the breast and pancreatic cancer ColX modules, as well as in their overlapping gene sets (Table S3A–C). Notably, GO categories including “cell migration” and “cell motility” were enriched in both cancer types, as were “ossification,” “cartilage development,” “skeletal system development,” and several terms related to Wnt signaling. Reactome pathways relating to collagen organization and ECM structure, function and degradation were similarly enriched, along with the role of *RUNX2* in regulating chondrocyte maturation (Table S3F–H).

In addition to these shared GO categories, breast and pancreatic cancer-specific ColX modules were individually enriched for several categories associated with the epithelial-to-mesenchymal transition (EMT). Enriched GO categories specific to the BRCA ColX module included “epithelial cell migration” and “mesenchymal cell proliferation,” and several Reactome pathways relating to signaling through FGFR2 were also significantly enriched (Table S3A, S3F). PAAD-specific GO hits included several developmental regulators such as Frizzled and Smoothened, which were corroborated by enrichment of Reactome pathways relating to Wnt, Hedgehog, and *RUNX2* signaling (Table S3B, S3G).

### ColX-associated gene networks are more strongly linked to OA and pro-metastatic processes in male pancreatic cancer cohorts

To assess the possible differential contribution of gender to the ColX-related pathology of pancreatic cancer, we also used WGCNA to define ColX modules in male and female patient subsets of the PAAD dataset, yielding 572 and 219 genes respectively, of which 144 were shared between male and female cohorts (Figure 2B). As male patients comprised only 1% of the BRCA dataset, we did not perform gender-specific analysis for breast cancer.

While both gender-specific pancreatic cancer modules were enriched for numerous GO terms and Reactome pathways which were also significant in their combined analysis (e.g., various ECM terms, “ossification,” “cartilage development,” and several terms related to chondroitin sulfate metabolism), the male PAAD ColX module was additionally enriched for the GO terms “bone morphogenesis,” “Wnt signaling,” and “epithelial to mesenchymal transition,” as well as Reactome pathways relating to *RUNX2* signaling and platelet responses (Table S3D, S3I). In contrast, the female PAAD ColX module was not significantly enriched for these terms or pathways; the primary hits were largely similar to those of the full PAAD ColX module (Table S3E, S3J), with inference of a more functionally-restricted ColX-associated genetic network possibly due to the smaller sample size.

### Breast and pancreatic cancer-specific ColX modules overlap with key matrisome and oncogenic pathway gene sets

To assess processes and gene functions captured within these four dataset-specific ColX modules, we performed statistical overlap analyses with multiple well-characterized gene sets encompassing a broad range of physiological and clinical functions. Both breast and pancreatic ColX modules were found to overlap significantly with matrisome gene sets, including ECM glycoproteins, collagens, ECM regulators, basement membranes, proteoglycans, and secreted factors^2^ (Figure 2C). The ColX modules include numerous matrisome genes previously implicated in tumor progression and metastasis, including ECM regulators *CTSB*, *LOXL2*, and *SERPINF1*, and ECM glycoprotein *SNED1*, which have been reported as markers of highly metastatic breast carcinomas that promote tumor invasiveness across a variety of models^68^ (Table S2A). *CTSB* is also upregulated in pancreatic cancer and may indicate increased activity of *CSTB*, which enhances later metastatic extravasation in PDAC; additionally, several collagens that are highly expressed in PDAC relative to its precursor pancreatic intraepithelial neoplasia (*COL6A1*, *COL6A2*, and *COL11A1*) are present in the pancreatic ColX modules^69,70^ (Table S2A). Thus, *COL10A1* is associated with common matrisome features that drive cancer progression, suggesting that ColX-associated genes may broadly play important roles in the development, maintenance, and pro-metastatic function of the tumor microenvironment^1,71^.

Multiple hallmark gene sets were significantly enriched in both breast and pancreatic ColX modules. The epithelial-to-mesenchymal transition (EMT) was the most significantly-overlapping hallmark gene set in both cancer types (p.adj = 5.2 × 10^-43^ and 3.2 × 10^-31^ for breast and pancreatic cancer, respectively; p.adj = 8.0 × 10^-40^ and 3.0 × 10^-17^ for male and female pancreatic cancer cohorts, respectively) (Figure 2D; Table S4). We observed substantial co-modularization of *COL10A1* with numerous collagen-binding integrin genes which play critical roles in invasion and blood vessel remodeling, including *ITGA1* (BRCA), *ITGA11* (BRCA, PAAD, and male PAAD), *ITGA5* (PAAD and male PAAD), *ITGAV* (BRCA), and *ITGB1* (BRCA and female PAAD) (Table S2A). The BRCA and male PAAD ColX modules additionally contain *DDR2*, a *COL10A1* receptor which has been shown to sustain the EMT phenotype to promote metastasis in breast cancer as well as induce EMT to accelerate the progression of pancreatic cancer^72,73^, consistent with fibrosis and metastasis-associated EMT pathway genes being most strongly enriched in those two ColX modules. Notable overlap was also observed across both cancer types with numerous other hallmark gene sets related to cancer cell motility and aberrant cell repair, including angiogenesis, coagulation, apical junction, myogenesis, and decreased UV repair response. The lists of EMT hallmark genes present in both breast and pancreatic ColX modules includes not only multiple collagen genes but also various genes coding for ECM proteins involved in non-collagenous network formation and cartilage and bone development (*ADAM12*, *COMP*, *MATN3*, *FN1*) (Figure S2E). Several of these genes are similarly present in both male and female pancreatic ColX modules, indicating conservation of pro-metastatic function across gender (Figure S2F). The pairwise concordance in EMT hallmark genes overlap between these groups is statistically significant by Fisher’s exact test (p = 6.2 × 10^-17^ between BRCA and PAAD; p = 4.6 × 10^-8^ between male and female PAAD), suggesting conservation of ColX module function across cancers and genders.

Several additional cancer type-specific hallmarks emerged as significant, including genes related to KRAS in breast cancer, and Notch signaling in pancreatic cancer. Interestingly, there were gender-specific differences within the pancreatic ColX modules, with the male ColX module significantly overlapping with genes involved in Hedgehog and Notch signaling, while neither of these gene sets overlapped significantly with the female ColX module (Figure 2D). In the female pancreatic cancer cohort, Hedgehog signaling was not significantly enriched in any module and Notch signaling was only enriched in module #12 (Table S4D), suggesting a stronger association between ColX expression and developmental pathways in male pancreatic tumors. Together, these observations provide evidence that ColX-associated gene signatures may confer more aggressive tumor features through activity of specific developmental pathways. Combined with the IHC observation that ColXα1 is expressed in tumor stroma (Figure 1B), these findings suggest that *COL10A1* expression is associated with an oncogenic fibroblast environment.

### ColX is associated with regulation of proliferative and mesenchymal cell states

To identify potential impacts on transcription factors (TFs), we assessed representation of transcription factor target (TFT) gene sets from the Gene Transcription Regulation Database (GTRD) in each ColX module. Two GTRD-annotated TFT gene sets were significantly enriched in both BRCA and PAAD ColX modules, corresponding to targets of *FOXH1* and *IGLV5-37* (Figure 2E). Corroborating the GO enrichment of EMT and related pro-metastatic pathways, *FOXH1* is an inducer of the TGF-β/Nodal/Activin signaling pathway and has been implicated in proliferation, migration, and invasion of both breast and pancreatic cancer^74,75^ (Table S5A). Of note, the latter TFT gene set appears to have been misattributed to *IGLV5-37* (an immunoglobulin with no known TF activity) in the GTRD database, and according to the source publication actually represents targets of the fusion oncoprotein *SS18-SSX*, which alters the normal regulatory activity of the SWI/SNF (BAF) ATP-dependent chromatin remodeling complex to drive oncogenesis in synovial sarcoma through induction of *SOX2*^76^. Both of these TFT gene sets include ColX module genes *ADAMTS6* (a metalloproteinase) and *RUNX1*, which drives mesenchymal stem cell proliferation and differentiation of myofibroblasts^77^; additionally, the *SS18-SSX* targets include *LRRC15*, a ColX module gene marking TGF-β-driven, myofibroblastic CAFs which may play an immunoregulatory role in PDAC^78^ (Table S5A). The shared enrichment of these two TFT gene sets in both ColX modules points toward a common association with fibroblast proliferation and invasion in breast and pancreatic cancer.

Among the BRCA ColX module genes, transcription targets of *PRMT5*, *SIX1*, *SUPT16H*, *PRDM4*, *TFEB*, *NFKBIA*, *ZNF585B*, and *PRDM5* were highly enriched (Figure 2E). These TFs have been established to regulate numerous pathways in breast and other cancers, ranging from cell proliferation and tumorigenesis to invasion and immune regulation (Table S5A). In particular, *SIX1* has been shown to induce EMT and metastasis in breast tumors^79^ and implicated in TGF-β regulation of collagen deposition leading to hampered immune infiltration and survival in cancer^80^, while *PRDM5* has been linked to pro-metastatic production of collagen in breast cancer^81^. The PAAD ColX module is enriched for targets of the transcription factor *NME2*, which regulates cancer cell proliferation in a variety of contexts through activation of *MYC* (Table S5A). Of note, *NME2* is an upstream regulator of *CTSK*, a protease primarily expressed in osteoclasts which is involved in ECM degradation and bone remodeling and has been found to be expressed in numerous cancers including breast carcinoma^82,83^.

### Increased ColX module expression is associated with activation of pro-metastatic pathways as well as immunosuppressive and myofibroblastic signatures

To investigate the physiological states associated with expression of ColX-related gene networks, we stratified patient samples into high and low ColX module expression groups and performed Quantitative Set Analysis for Gene Expression (QuSAGE) to identify differentially-expressed pathways between these cohorts (see Methods section and Figure S3 for definition and derivation of %G.A.M.E. metric). QuSAGE is an efficient alternative to classical Gene Set Enrichment Analysis (GSEA) which accounts for inter-gene correlations and provides a more robust quantification of differential pathway expression by generating a complete probability density to describe the activity of a particular gene set of interest^61–63^. We ran QuSAGE on ColX module-stratified samples for the MSigDB hallmark pathways^54^, a collection of previously-characterized immune signature pathways^59^, the MSigDB GTRD TFT gene sets^54,55^, and a collection of CAF genesets curated from human breast and pancreatic cancer datasets^19^.

Increased activity of hallmark pathways associated with aggressive cancer phenotypes was significantly associated with ColX module expression (Figure 3A). Heightened ColX module expression was positively associated with activation of genes involved in EMT (2.5-fold average increase across all cohorts), angiogenesis (1.7-fold average increase), coagulation (1.6-fold increase in BRCA; also significant in male and female PAAD cohorts), and myogenesis, apical junction and apical surface (each 1.5-fold increase in male PAAD cohort). Additionally, QuSAGE revealed increased activity of genes typically decreased in response to UV exposure across all cohorts (1.5-fold average increase). We also observed significant upregulation in high-ColX module samples of several regulatory pathways involved in tumor progression and immune response that were not identified through gene enrichment testing, including inflammatory response and KRAS signaling (all cohorts), Hedgehog signaling (BRCA), allograft rejection, IL-6/JAK/STAT3 signaling, and TNF-α signaling via NFKB (all PAAD cohorts), IFN-γ response (PAAD and male PAAD cohorts), complement (male and female PAAD cohorts), and IL-2/STAT5 signaling and TGF-β signaling pathways (male PAAD cohort). Several hallmark gene sets related to proliferation were downregulated in high-ColX module BRCA samples, including oxidative phosphorylation, MYC targets (V1, V2), E2F targets, and G2/M checkpoint; the high-ColX module male PAAD cohort also exhibited downregulation of MYC targets (V2) and oxidative phosphorylation. These findings indicate that tumors with high *COL10A1* expression exhibit increased activity of numerous pathways related to pro-metastatic and inflammatory processes, along with a concomitant decrease in activity of homeostatic pathways such as oxidative metabolism and cell cycle regulation. Thus, activity of ColX module genes highlights more aggressive cancer phenotypes.

**Figure 3:**
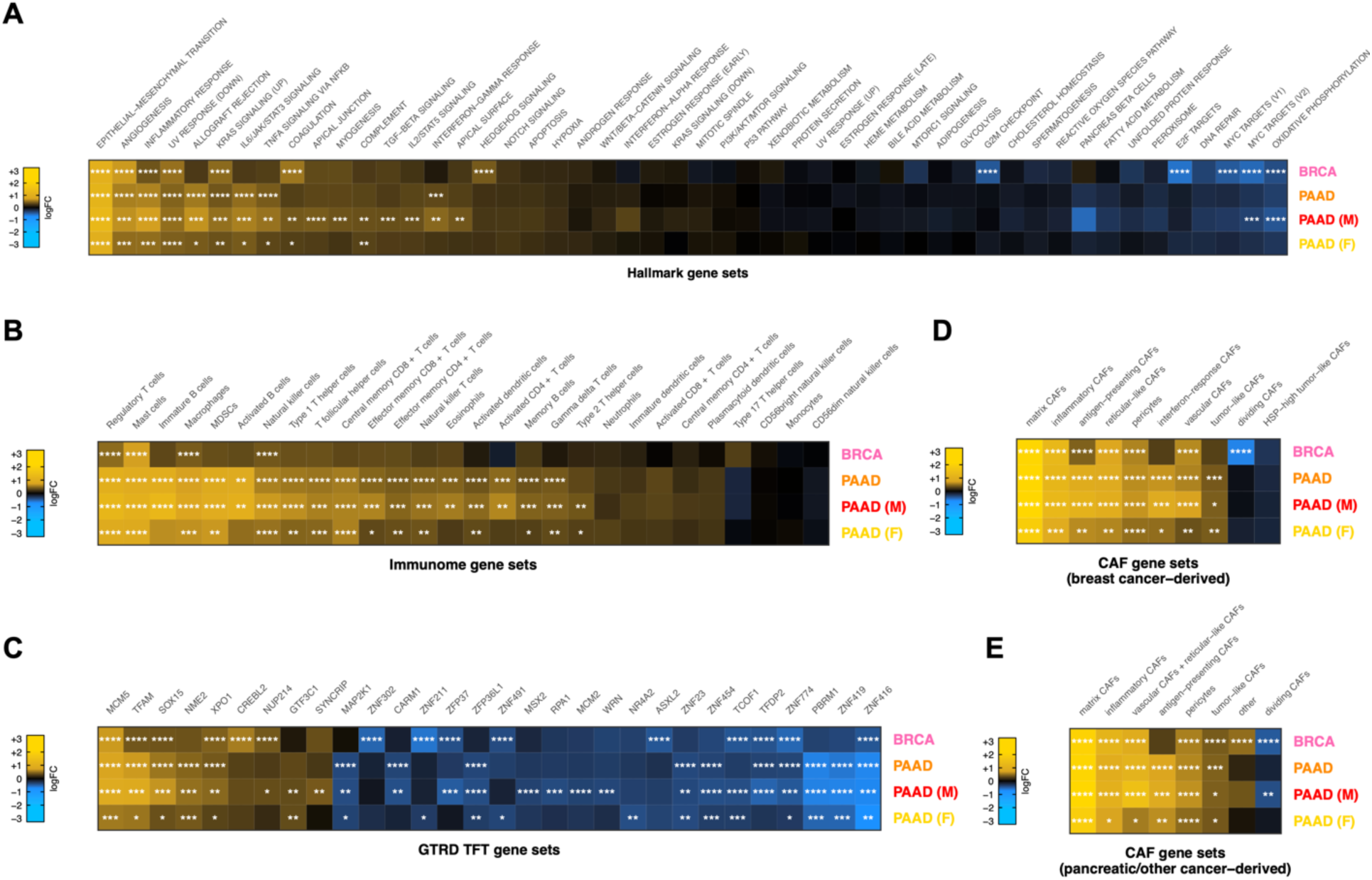
Tumors with high ColX module expression exhibit activation of pro-metastatic, immunosuppressive, and myofibroblastic gene signatures. (A–E) Mean pathway activation (by QuSAGE) of **(A)** MSigDB hallmark pathway gene sets^54^, **(B)** cancer immunome gene sets^59^, **(C)** top significant GTRD transcription factor target gene sets^55^, and cancer-associated fibroblast (CAF) gene sets derived from **(D)** breast tumors and **(E)** pancreas and other tumors^19^, in samples with high ColX module expression relative to samples with low ColX module expression for each cancer dataset. Gene sets within each block are ordered by mean pathway activation rank across all 4 datasets. See Figure S4 for QuSAGE analysis of all modules and Table S5B for reported functions of selected enriched TFTs. Significance values: *, p.adj < 0.05; **, p.adj < 0.01; ***, p.adj < 0.001; ****, p.adj < 0.0001.

ColX module expression was also significantly associated with increased expression of numerous immune signatures in both BRCA and PAAD, including immunosuppressive and tumor-promoting features such as regulatory T cells, mast cells, and tumor-associated macrophages (TAMs) (Figure 3B). In pancreatic cancer specifically, we observed broader upregulation of numerous immune cell types, but notably no upregulation of the activated CD8^+^ T cell signature, and greater upregulation of immature vs. activated B-cell signatures. Although the activation of regulatory T cells was not uniquely associated with ColX module expression, we note that other WGCNA modules tracking positively with regulatory T cell activity were almost invariably also associated with activated CD8^+^ T cell signal as expected in controlled physiological immune responses, while the ColX modules were not (Figure S4A–D). ColX module expression was consistently associated with regulatory T cell activity even in the absence of activated CD8^+^ T cells. These findings suggest that the ColX modules are strongly associated with immunosuppressive environments.

QuSAGE analysis of GTRD TFT gene sets highlighted numerous transcription factors whose downstream targets were differentially expressed in tumors with high ColX module expression, several of which are known to modulate tumor advantage in breast or pancreatic cancer (Figure 3C; see Table S5B for detailed descriptions of significant TFs). Targets of *MCM5*, a crucial component of the MCM replicative helicase complex which has been linked to negative prognosis in breast cancer and implicated as a marker of pancreatic malignancy, were the most significantly upregulated in both BRCA and PAAD high-ColX module tumors, an effect which was also observed in the pancreatic gender-specific analysis. Expression of *NME2* targets was also increased significantly in high-ColX module PAAD tumors; this protein is a well-established activator of *MYC* and while competing evidence suggests that it may have pro- or anti-tumor effects in diverse contexts, it has been identified as upregulated in metastatic PDAC samples compared to primary tumors by snRNA-Seq^84^. *XPO1* and *NUP214*, two associated proteins whose regulatory targets are mutually upregulated in high-ColX module tumors, have collectively been described as drivers of breast cancer and markers of poor survival in pancreatic cancer. *NUP214* is of particular interest as its varied fusion products have been implicated as drivers of breast cancer (via fusion with *NOTCH*)^85^ and leukemogenesis (via aberrant activation of HOX genes)^86^; similarly, *HOXC8* co-modularizes with *COL10A1* in BRCA (Table S2A) and *Hoxa3* has been shown to be upregulated alongside *Col10a1* and implicated in OA progression in hypertrophic mouse chondrocytes^87^. Targets of *GTF3C1* and *SYNCRIP*, both of which are upregulated in metastatic pancreatic tumors compared to primary tumors, were also significantly increased in expression in high-ColX module gender-specific PAAD cohorts.

Conversely, targets of multiple key genes crucial to the maintenance of various tumor-suppressive protein complexes were significantly downregulated in high-ColX module samples (Figure 3C), including *ASXL2* in BRCA and *PBRM1* in PAAD (irrespective of gender). Significant downregulation in high-ColX module tumors was also observed for targets of numerous zinc-finger TFs (*ZNF302*, *ZNF211*, *ZFP37*, *ZNF23*, *ZNF454*, *ZNF774*, *ZNF419*, and *ZNF416*) associated with *TRIM28*, a pro-tumorigenic driver of EMT which stabilizes *TWIST1* to promote cancer cell invasion and migration^88^. While individual coexpression of each significant zinc-finger TF with *TRIM28* and *TWIST1* varied across cohorts, on average they were negatively correlated with expression of both pro-metastatic genes in BRCA and PAAD, potentially corroborating the increased activity of EMT drivers in high-ColX module tumors.

ColX module expression was additionally associated with increased activity of multiple CAF gene sets. We analyzed gene signatures from diverse CAF populations recently characterized by Cords et al. using scRNA-Seq data from breast tumors (∼14,000 CAFs from 14 patients with breast invasive carcinoma) and pancreatic tumors (∼5,700 CAFs from 4 cancer types, of which 30% were sourced from patients with pancreatic ductal adenocarcinoma)^19^. As these gene signatures have been shown to be highly conserved across cancer types, we performed matched and cross-comparisons between breast- and pancreatic-derived CAF signatures and our breast and pancreatic cancer data, stratified by ColX module expression. We found that gene sets associated with numerous breast-derived myofibroblastic CAF populations (matrix, inflammatory, and antigen-presenting) were significantly upregulated in high-ColX module samples compared to low-ColX module samples in both BRCA and PAAD cohorts, as were genes associated with vascular CAFs, which have been shown to facilitate angiogenesis and tumor vascularization (Figure 3D). High ColX module expression in the PAAD cohorts was uniformly associated with increased interferon-response and tumor-like CAF signatures as well. Matrix CAFs (mCAFs) play a role in ECM remodeling, migration, TGF-β-driven myofibroblastic activation, and EMT; numerous mCAF markers are present in the ColX modules, including *COMP*, *MMP11*, *POSTN*, *COL1A1*, *COL1A2*, *LRRC15*, and the pro-myofibroblastic markers *FAP* and *PDPN* (Table S2A). Various other CAF-specific genes are co-modularized with *COL10A1*, including inflammatory CAF markers *CXCL12* (BRCA) and *CXCL14* (female PAAD), vascular CAF markers *ACTA2* (all ColX modules) and *NOTCH3* (PAAD and male PAAD), and tumor-like CAF markers *PDPN* (BRCA, PAAD, and male PAAD) and *TMEM158* (male PAAD). Additionally, the dividing CAF signature was significantly decreased in high-ColX module BRCA tumors, corroborating the decrease in homeostatic cell cycle control indicated by the hallmark pathway analysis of the BRCA cohort (Figure 3D). Similar enrichment of pancreatic-derived myofibroblastic CAF signatures in high-ColX module tumors was observed in both BRCA and PAAD cohorts, concomitant with increased signal associated with tumor-like CAFs and decreased signal associated with dividing CAFs (Figure 3E). These observations further support the theme of ColX being a marker of CAF heterogeneity in stromal environments across diverse cancers.

### ColX gene and module expression levels are prognostic of differential survival outcomes

Given the relevance of the stromal microenvironment to cancer progression and patient outcomes, we assessed the predictive value of ColX with regard to breast and pancreatic survival metrics. Using a multivariate Cox proportional hazards model to assess overall survival risk, increased *COL10A1* expression was found to be significantly associated with negative prognosis (+17% risk per standard deviation in breast cancer, p.adj = 0.048; +31% risk per standard deviation in pancreatic cancer, p.adj = 0.030) (Figure 4A). These hazard contributions were conferred in addition to the significant effects of age (+4% risk per year in breast cancer, p.adj = 2.2 × 10^-8^; +3% risk per year in pancreatic cancer, p.adj = 0.030) and advanced tumor stage (+182% risk for stages 3-4 compared to stages 1-2 in breast cancer, p.adj = 7.3 × 10^-9^).

**Figure 4:**
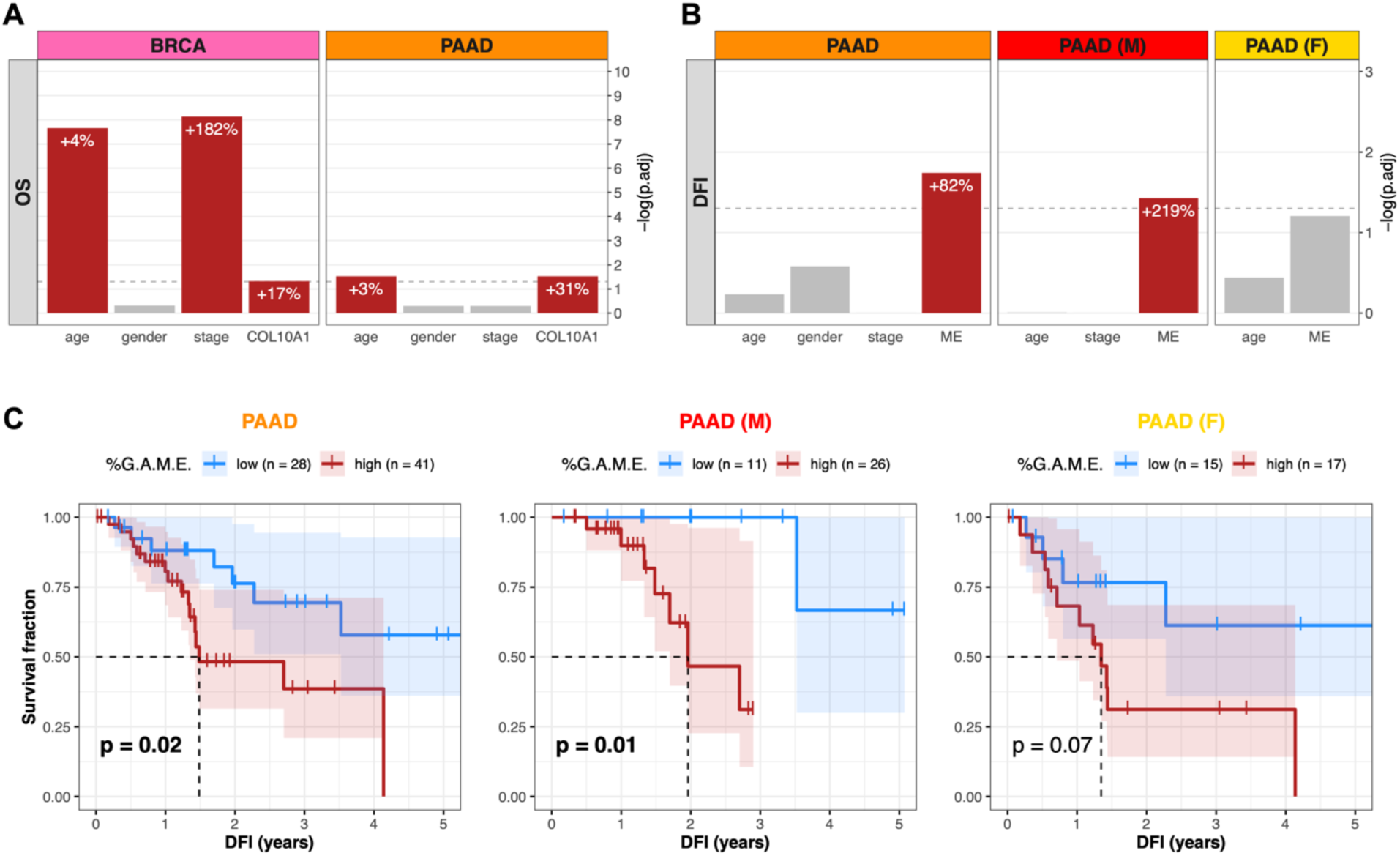
*COL10A1* and ColX module expression stratify breast and pancreatic cancer cohorts by survival outcomes. **(A and B)** BH-adjusted significance values for multivariate Cox proportional hazards models conditioning either **(A)** overall survival (OS) on age, gender, binarized tumor stage, and *COL10A1* gene expression, or **(B)** disease-free interval (DFI) on age, gender, binarized tumor stage, and ColX module eigengene expression (ME). Dotted lines correspond to p.adj = 0.05. Per-unit contributions of each significant variable (red bars) to the overall hazard risk in each panel is indicated by white percentages (for *COL10A1* and ColX ME expression, 1 standard deviation = 1 unit). Note that the Cox model for the female pancreatic cancer cohort was not conditioned on binarized tumor stage because all samples were classified into the same group. See Figure S5 for full survival analysis results for all WGCNA-inferred modules. **(C)** DFI Kaplan-Meier survival curves for the pancreatic cancer cohorts based on %G.A.M.E. groupings. Dotted lines correspond to median “survival” (i.e., time to recurrence). Shaded regions represent the 95% confidence intervals for each group. Log-rank test p-values are shown for each panel.

To determine whether ColX-associated gene modules preserved the prognostic value of ColX itself, we performed a similar analysis using the ColX module eigengene (ME) in place of *COL10A1* expression. Increased ColX module expression was significantly associated with shortened disease-free interval (DFI) in pancreatic cancer (+82% risk per standard deviation, p.adj = 0.018) (Figure 4B). This effect was also significant in the male pancreatic cancer subcohort (+219% risk per standard deviation, p.adj = 0.037), but was not preserved in the female subcohort (p.adj = 0.062). The ColX ME did not significantly impact DFI in breast cancer, possibly due to the greater availability and efficacy of curative therapies for breast cancer relative to pancreatic cancer. However, the fact that the ColX ME was one of only two modules in the pancreatic cancer cohort (as well as in the male subset) whose expression tracked significantly with increased DFI risk suggests that the ColX signature is uniquely associated with likelihood of recurrence of advanced cancer (Figures S5B, S5C).

We then investigated whether the rescaled metric of ColX module expression (%G.A.M.E.) could be used to effectively stratify pancreatic cancer patients based on DFI. Patients with high %G.A.M.E. had significantly worse DFI prognosis compared to those with low %G.A.M.E. (p = 0.02) by Kaplan-Meier analysis. Interestingly, this effect was preserved in the male subcohort (p = 0.01), but attenuated in the female subcohort (p = 0.07) (Figure 4C). These findings recapitulated the multivariate hazard risk analysis, corroborating the prognostic value of ColX expression at both the gene and module level.

### ColX module expression correlates with increased activity of bone marrow stroma and osteoarthritic cartilage signatures

Identifying connections between the roles of ColX in cancer and non-cancer cells may provide novel insights into common factors at play in these tissues. In its canonical contexts, *COL10A1* is a marker for hypertrophic chondrocytes and bone marrow-derived mesenchymal stem cells and contributes to cellular senescence, ECM degradation, and angiogenic invasion during OA^6,9,11,89–92^. Therefore, we examined the bone marrow and OA character of breast and pancreatic tumors based on their levels of ColX module expression, using RNA-Seq data from four types of cartilage or bone marrow stromal cells for comparison^10^. While *COL10A1* was not strongly expressed by normal cartilage stromal cells (NCSCs), it was significantly upregulated in both bone marrow stromal cells (BMSCs; over 100-fold increase, p.adj = 1.1 × 10^-31^) and OA mesenchymal stromal cells (OA-MSCs; 18.6-fold increase, p.adj = 3.4 × 10^-11^) (Figure 5A).

**Figure 5:**
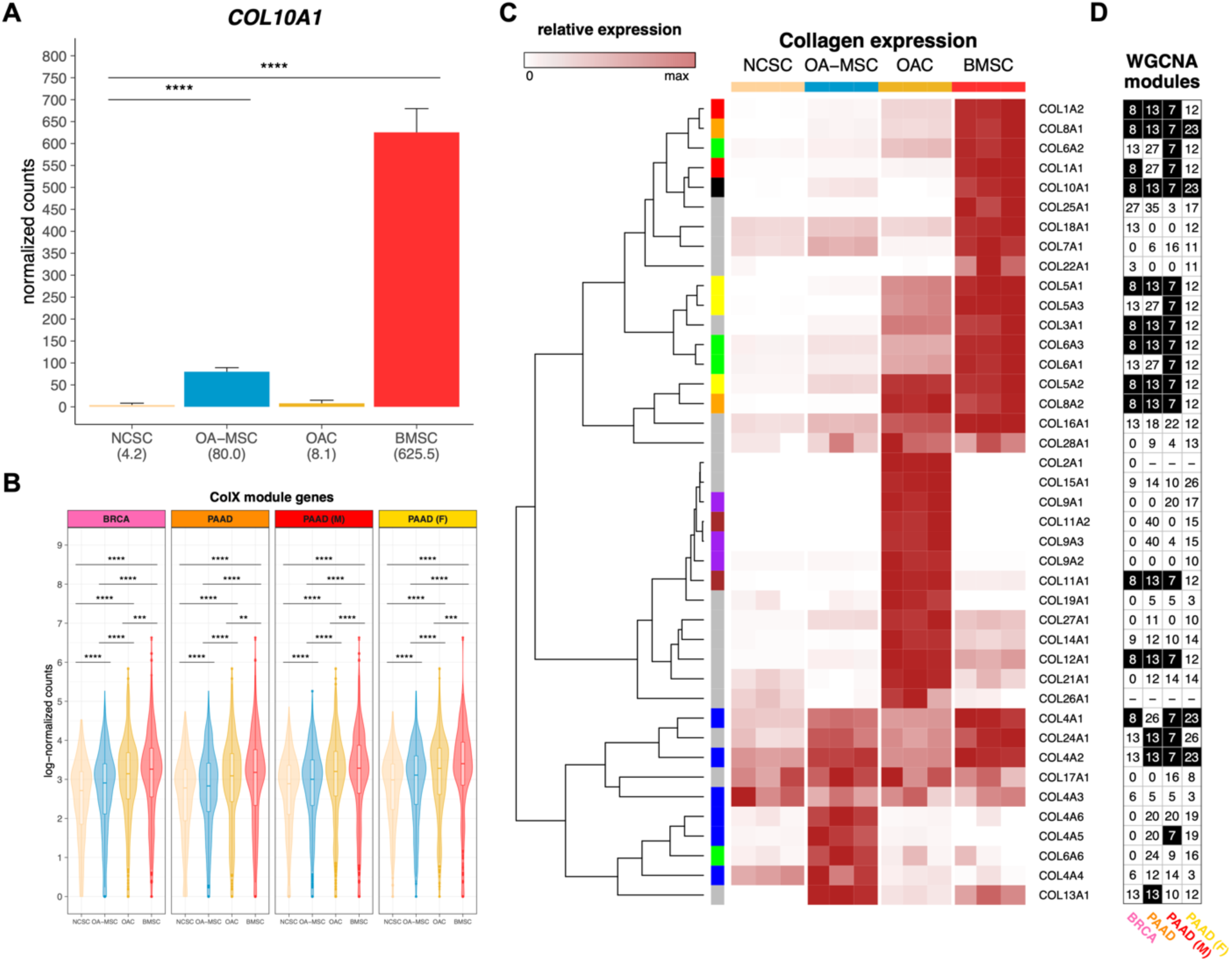
Bone marrow and cartilage cells are differentiated by collagen expression clusters. **(A)** Normalized expression of *COL10A1* across bone marrow and articular cartilage cell types (*n* = 3 each). Mean normalized expressions are indicated in parentheses. Significance values were computed using DESeq2, BH-adjusted across all genes analyzed. **(B)** Log-normalized expression of ColX module genes in each cell type (*n* = 3 each). Normalized gene-wise expressions were averaged prior to log-transformation with pseudocount of 1. Significance values were computed using the paired Wilcoxon signed-rank test on log-normalized counts. **(C)** Relative expression of collagen genes across cell samples (*n* = 3 each). Normalized expression values are scaled by rows. *COL10A1* is indicated by a black box in the left column; genes contributing to the same parent collagen are indicated by same-color boxes; and genes which are the sole contributor to their parent collagen are indicated by gray boxes. Note that *COL6A4P1*, *COL6A5*, *COL20A1*, and *COL23A1* were filtered out as “low-expression” genes. See Figure S7A for normalized (unscaled) expression of all collagen genes across cell types. **(D)** WGCNA module assignments for all collagens in each cancer dataset. See Table S2B for WGCNA module assignments for all genes. Black cells indicate the ColX module for each column. “-” indicates low-expression genes which were filtered out; genes labeled “0” were not assigned to any module by WGCNA. See Figure S7B for normalized expression of all collagen genes across TCGA cohorts.

Interestingly, the module-wide expression of *COL10A1*-associated genes in each cancer was significantly increased in BMSCs, followed by two OA cartilage cell types, OA chondrocytes (OACs) and OA-MSCs (Figure 5B); additionally, cell type-specific markers for BMSCs and OACs were found to be highly enriched for genes in the ColX modules (Figure S6A–D). These enrichments were significant by Fisher’s exact test when compared across all WGCNA modules for both BRCA (p.adj = 1.6 × 10^-15^ for BMSC and p.adj = 4.7 × 10^-3^ for OAC) and PAAD (p.adj = 1.7 × 10^-11^ for BMSC and p.adj = 0.02 for OAC) (Figure S6E). Comparative analysis of the expression of various collagens across cartilage and bone marrow cell types revealed that the collagen gene signatures of BMSCs were closely aligned with the ColX modules in BRCA and PAAD, in particular male but not female PAAD (Figures 5C, 5D). Numerous BMSC-specific collagens also co-modularized with *COL10A1* in tumors, including *COL8A1* and *COL8A2*, *COL1A1* and *COL1A2*, *COL4A1* and *COL4A2*, and several genes contributing to collagen types III, V and VI. These latter three collagens have all been implicated in tissue repair, wound healing, expression profiles of diverse CAF subtypes, and regulation of both BMSC and OAC physiology^6,10,93^. Like *COL10A1*, collagen type IV is a network-forming collagen; its α112 triple helical form is ubiquitously expressed in basement membranes and is present in both normal and OA articular cartilage, where it is expressed by chondrocytes^6,94^. *COL8A1* is highly expressed in OA tissues and, like *COL10A1*, is a myofibroblastic-specific marker in cancer^93,95^. These results suggest that, in addition to *COL10A1*, BRCA, PAAD and male PAAD tumors especially share common ECM gene signatures with BMSCs.

Cell type proportion inference using CIBERTSORTx revealed that expression of ColX modules was positively associated with signatures attributable to BMSCs, OACs, and OA-MSCs across both breast and pancreatic tumors (Figures 6A, 6B). ColX module expression was significantly and positively correlated with CIBERSORTx-inferred absolute proportions of BMSCs as well as, to a lesser extent, two OA cell types (OACs and OA-MSCs) (Figure 6C). Conversely, ColX module expression was negatively correlated with the absolute inferred proportion of NCSCs in breast cancer; correlations of similar direction and magnitude were observed in pancreatic cancer, with some loss of significance likely attributable to decreased power due to the smaller cohort size (Figure 6C). The relative inferred proportion of NCSCs among the 4 cell types also decreased significantly with increased ColX module expression in both cancers, concomitant with a significant increase in the relative inferred proportion of BMSCs (Figure 6D). Thus, ColX emerged as a biomarker for bone marrow-derived cells in BRCA and PAAD tumors.

**Figure 6:**
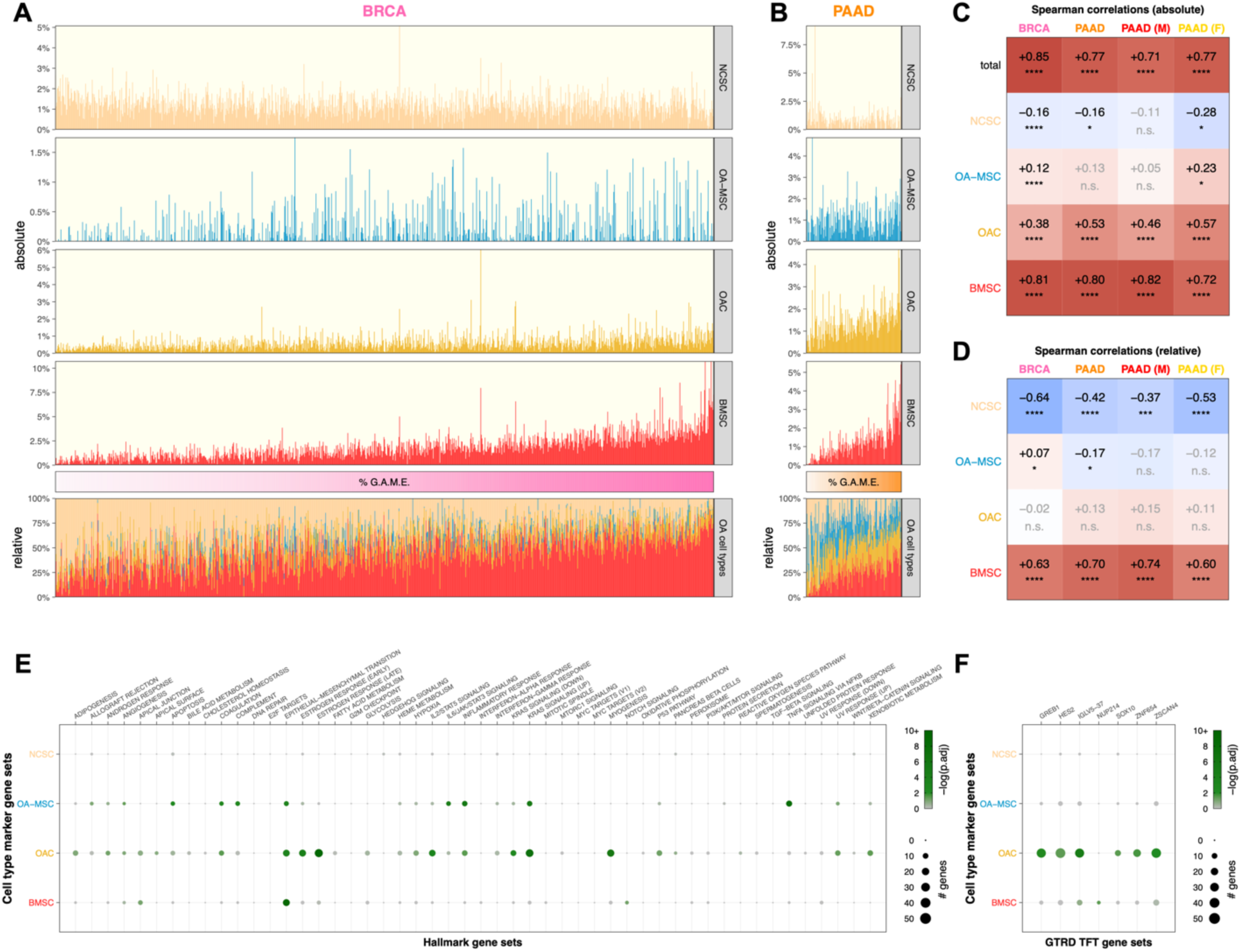
Tumoral proportions of osteoarthritic cell types trend with ColX module expression and cancer-associated pathway activity. **(A and B)** Sample-wise absolute (*top*) and relative (*bottom*) proportions of 4 bone marrow and cartilage cell types inferred by CIBERSORTx for TCGA **(A)** breast and **(B)** pancreatic cancer cohorts. Color bars (*middle*) indicate relative cancer-specific ColX module expression; samples were ordered by increasing %G.A.M.E. See Methods section and Figure S3 for details on %G.A.M.E. metric. **(C and D)** Spearman correlations between relative cancer-specific ColX module expression and **(C)** absolute or **(D)** relative OA cell type proportions inferred by CIBERSORTx. Raw p-values were Bonferroni-corrected across rows. Significance values: *, p < 0.05; **, p < 0.01; ***, p < 0.001; ****, p < 0.0001; n.s., not significant. NCSC, normal cartilage stromal cells; OA-MSC, osteoarthritis mesenchymal stromal cells; OAC, osteoarthritis chondrocytes; BMSC, bone marrow stromal cells. **(E and F)** Bubble plots of **(E)** MSigDB hallmark pathway gene sets^54^ and **(F)** top significant GTRD transcription factor target gene sets^55^ in cell type-specific marker gene sets. Of note, the TFT gene set attributed to *IGLV5-37* in the GTRD database actually represents targets of the fusion oncoprotein *SS18-SSX*, as described in the text. See Methods section for definition of bone marrow and cartilage cell type-specific marker genes. Significance values: *, p.adj < 0.05; **, p.adj < 0.01; ***, p.adj < 0.001; ****, p.adj < 0.0001.

### ColX highlights similar EMT markers between OA disease states and tumors

EMT comprises a range of markers depending on tissue settings and surrounding phenotypes, and contributes to diverse processes including gastrulation, fibroblastic differentiation, and tumor metastasis^96^. Examination of cell type-specific markers for each ColX-expressing non-cancer cell type (Table S6) revealed that EMT hallmark pathway genes were significantly overrepresented in BMSCs, OACs, and OA-MSCs (Figure 6E). Notably, several EMT-associated genes were identified as both OA cartilage cell type markers and ColX module genes. *COL11A1*, *COMP*, and *MATN3*, which were present in both breast and pancreatic ColX modules (Table S2A; Figures S2E, S2F), are markers of chondrogenesis expressed distinctly by OACs (Table S6); additionally, *COMP* is a potent anti-apoptotic factor which may contribute to survival of both chondrocytes and neoplastic cells. The BRCA ColX module also includes the OAC-specific markers *LUM* (also present in the PAAD ColX module) and its related genes *DCN* and *ECM2*, which are collectively involved in collagen fibril organization and epithelial cell migration (Table S2A). Genes coding for various CXC chemokines, inflammatory cytokines, and matrix metalloproteinases were identified as EMT-associated markers in OA-MSCs (*CXCL1*, *CXCL6*, *CXCL8*, *IL6*, *MMP1*, *PTX3*, *TNFAIP3*) (Table S6); while these were not specifically co-modularized with *COL10A1* in tumors, several related genes in both the CXC and MMP families are present in the ColX modules (pro-metastatic *CXCL12* in BRCA, pro-angiogenic *MMP2* in both BRCA and PAAD, and related MMPs 2, 3, and 14 in male PAAD) (Table S2A; Figures S2E, S2F).

Significant overlap in EMT pathway genes was observed between BMSC-specific markers and the ColX modules; 14 BMSC markers co-modularized with *COL10A1* in BRCA (p = 1.8 × 10^-5^ by Fisher’s exact test), 10 of which also co-modularized in PAAD (p = 2.8 × 10^-3^ by Fisher’s exact test) (Figure S6F). Several of these overlapping EMT pathway genes are also highly expressed in matrix CAFs (*COL1A2*, *LRRC15*, *POSTN*); other overlapping genes have been shown to contribute to vascular remodeling (*ACTA2*, *CTHRC1*) and myofibroblastic motility (*CALD1* and *TPM1*, both of which are significantly upregulated in metastatic pancreatic cancer compared to primary tumors^84^) (Table S2A; Table S6; Figures S2E, S2F). In particular, *FN1*, *VCAN*, and *LRRC15* are markers of BMSCs which also co-modularized with *COL10A1* in cancer; all three genes are involved in cell migration, and *FN1* contributes directly to osteoblast mineralization as well as metastasis. Together, these results suggest a shared activation of common EMT-related pathways in cancer, bone marrow, and pathological OA contexts, highlighting the role of *COL10A1* as a marker of both BMSCs and pro-metastatic tumors. The ColX modules thus highlight well-characterized inflammatory OA-like disease states which contribute to a pro-metastatic tumor microenvironment in both breast and pancreatic cancer.

Several TFs whose targets were found to be differentially expressed in high-ColX module vs. low-ColX module tumors have also been reported to play roles in OA disease states (Figure 3C). Upregulated TFT sets included targets of *MCM5*, whose expression is increased in rat chondrocytes following inflammatory stimulation by IL-1β; *TFAM*, which promotes mitochondrial biogenesis in OA chondrocytes; and *NME2*, which has been shown to be upregulated in early OA (Table S5B). Downregulated TFT sets included targets of the associated proteins *RPA1*, whose expression is decreased in OA patients, and *WRN*, which is mutated in Werner’s syndrome and confers early-onset OA risk; as well as *PBRM1*, which has been implicated in GWAS of OA and regulation of BMP signaling and osteogenic fate determination (Table S5B). Cell type-specific markers for both OACs and OA-MSCs were also significantly enriched for various inflammatory/immune (e.g., interleukin/STAT protein signaling) and proliferative (e.g., KRAS signaling) pathways (Figure 6E). Additionally, OAC- and BMSC-specific markers were found to be enriched for targets of the TFs *SS18-SSX* (whose targets were misattributed in the GTRD database to the immunoglobulin *IGLV5-37* as noted above), and *NUP214*, respectively (Figure 6F). These findings corroborate the similar enrichment of *SS18-SSX* targets in both the breast and pancreatic ColX modules (Figure 2E), suggesting that common pathways involving dysregulated cell proliferation and tissue remodeling are involved in both cancer and OA (Table S5A), as well as the differential activation of *NUP214* targets in high-ColX module breast and male pancreatic tumor samples (Figure 3C), which additionally suggest a shared pathogenic mechanism linked to aberrant transcription regulation (Table S5B). Many TFs thus appear to be perturbed similarly in both high-ColX module tumors and OA development, suggesting similar dysregulation of transcriptional programs across these disease states.

## DISCUSSION

The pathological contributions of collagen type X (*COL10A1*, ColX) and its associated gene networks are especially notable in breast and pancreatic tumors, where they contribute to fostering an inflammatory and immunosuppressive microenvironment that contributes to aggressive tumorigenesis, metastasis, and poor clinical prognosis. A significant component of this pathological role appears to be related to ColX’s identity as a marker for bone marrow-derived cells and cancer-associated fibroblasts (CAFs), both of which are strongly associated with extracellular matrix (ECM) remodeling, the epithelial-to-mesenchymal transition (EMT), and progression of disease in both cancer and OA. The findings of this study, summarized in a graphical abstract for visualization (Figure 7), suggest that there is a close relationship between these two ColX-expressing stromal cell populations; one in breast and pancreatic cancer and the other in bone marrow and OA cartilage.

**Figure 7:**
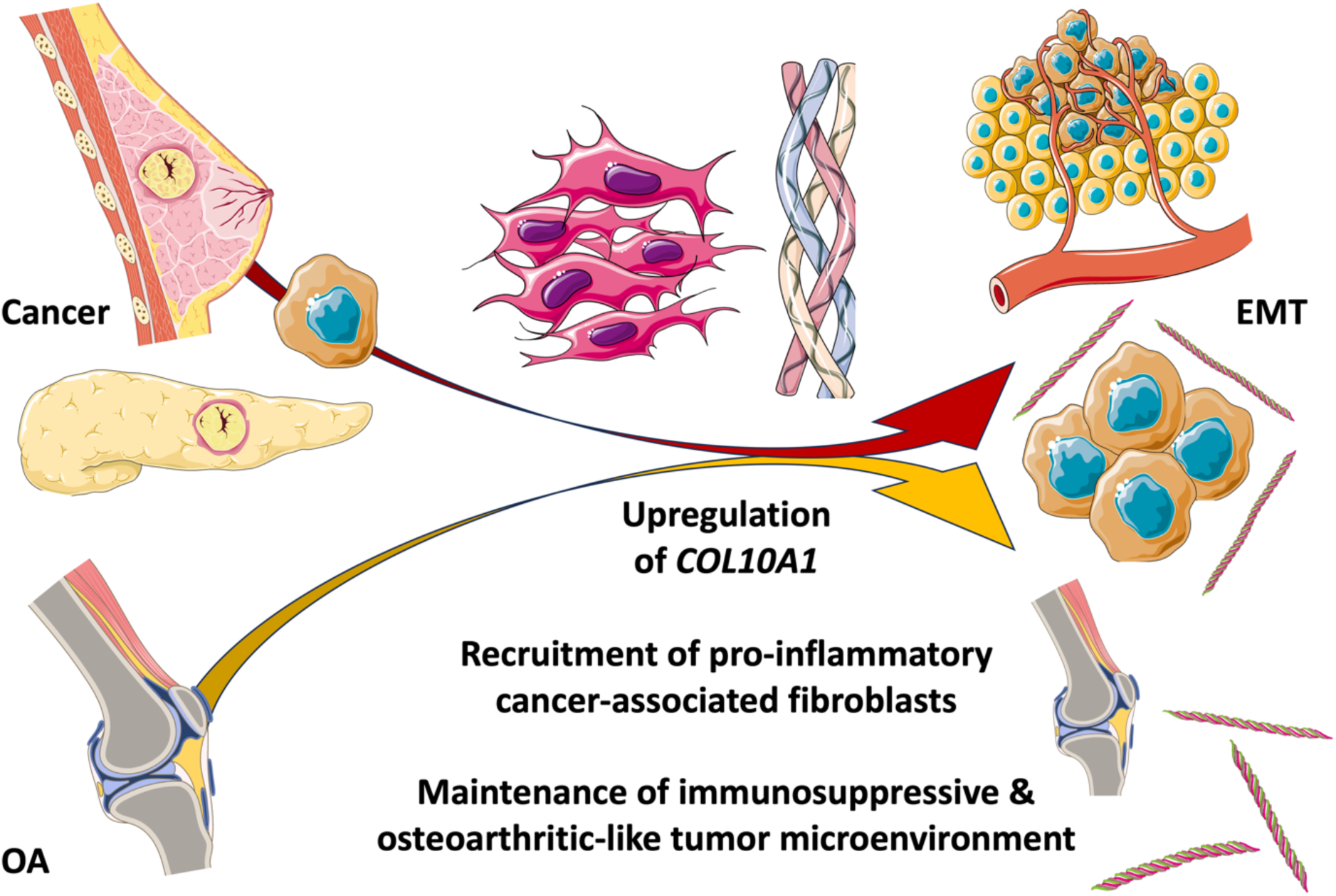
Pathological expression of *COL10A1* fosters immunosuppressive, fibroblastic microenvironments in cancer, bone marrow, and cartilage. Graphical abstract summarizing the findings presented in this study. In brief, a specifically expressed collagen, *COL10A1*, connects the ECM and tissue microenvironments across cancer and bone. The pathological contributions of ColX and its associated gene networks are especially prominent in breast and pancreatic tumors, and mimic the development of a similar inflammatory and fibroblast-dominated environment seen in bone marrow and cartilage changes in OA. A central outcome of this shared impact is the epithelial-to-mesenchymal transition (EMT), which contributes to disease progression in both contexts. *All images were sourced from Bioicons (*https://bioicons.com*) and are licensed for public use by Servier (*https://smart.servier.com*) under CC-BY 3.0 (*https://creativecommons.org/licenses/by/3.0*) or by DBCLS (*https://togotv.dbcls.jp/en/pics.html*) under CC-BY 4.0 (*https://creativecommons.org/licenses/by/4.0).

ColX is normally expressed only during skeletal development at the cartilage-to-bone transition and in bone marrow during adulthood. However, ColX is also highly expressed in certain disease contexts including OA cartilage and a variety of solid tumors, especially breast and pancreatic cancer. Here, we demonstrated that ColX expression is aberrantly elevated in solid tumors from TCGA; namely, breast invasive carcinoma (BRCA) and pancreatic adenocarcinoma (PAAD). From RNA-Seq analysis, we found that ColX is also highly expressed by bone marrow stromal cells (BMSCs) and senescent mesenchymal stromal cells (OA-MSCs) in OA. Recent work demonstrates that chromatin accessibility of *COL10A1* is highest in hypertrophic chondrocytes and increases with progression of chondrocyte maturation in skeletal development^97^. Indeed, in BRCA tumors we observed that *COL10A1* expression is anticorrelated with the somitic mesoderm marker *TCF15* (Spearman’s *ρ* = −0.16, p = 4.5 × 10^-8^) and positively correlated with the sclerotome marker *PAX9* (Spearman’s *ρ* = +0.31, p = 2.0 × 10^-26^), supporting its contribution to the development of paraxial mesoderm and ossification in cancer. ColX’s patterns of co-expression and variability across breast and pancreatic tumors suggest that it may be associated with more aggressive disease as indicated by activation of EMT and angiogenic pathways as well as heightened relapse and survival risk.

There are compelling reasons to extrapolate from the roles that *COL10A1* plays in OA to putative functions within the tumor microenvironment. ColX is known to contribute to calcified zones of articular cartilage when expressed by hypertrophic chondrocytes in OA, a phenotype similar to mechanical stiffness conferred by the stroma of many solid tumors^6, 98^. The EMT process is also enriched during the progression of OA; we have previously described this “chondrocyte-mesenchymal transition” (CMT) and note that it mimics the loss of tissue-specific gene expression and inflammatory increase in invasive capacity which is the hallmark of the metastatic process^10^. Invasion and aggression of breast tumors are known to be correlated with mechanical stress and stiffened phenotypes^99,100^, and pancreatic tumors have similarly been associated with such stress owing to their significant stromal fraction^101,102^. The shared nature of these phenotypes highlights the role of the ECM and stroma in tumor malignancy, and suggests that ColX may contribute in a similar pathological manner to both OA and cancer progression.

Numerous biological pathways and transcriptional networks implicated in pro-OA and pro-metastatic disease states are significantly enriched within or activated in conjunction with increased expression of the ColX modules characterized here. The hallmark pathway most significantly enriched in each ColX module was the epithelial-to-mesenchymal transition (EMT), which was also found to be differentially activated in high-ColX module expression samples compared to low-ColX module expression samples. EMT is intimately involved in the intercellular remodeling processes of embryonic development, fibrosis, and wound healing, and its activity in cancer is broadly considered crucial to the invasion of ECM and neighboring tissues which initiates metastasis in epithelial malignancies^103^. Upregulation of EMT was found to coincide with increased activity of angiogenic pathways which facilitate tumor cell intravasation and motility into the circulatory system, as well as key immunosuppressive cellular signatures (regulatory T cells and TAMs) which permit tumor development. Activity of both of these immune cell types have been associated with poor outcomes in breast and pancreatic cancer^104–107^. These results suggest that heightened ColX module activity signifies aggressive tumor states marked by impaired immune response and greater potential to metastasize.

ColX modules also exhibited significant overrepresentation and/or differential activity of transcription factor (TF) networks crucial to numerous developmental pathways known to play a role in cancer aggressiveness and metastasis (Notch, Wnt, and Hedgehog signaling), including *FOXH1*, *SOX15*, and several *PRDM* gene family members^108–113^. GO enrichment of each ColX module revealed multiple genes implicated in ossification, such as *RUNX2* and *TGFB3*, both of which have been shown to play a role in OA, influence developmental pathways such as Wnt signaling, and contribute to EMT in breast and pancreatic cancer^114–116^. The male PAAD ColX module was additionally enriched for GO terms relating to EMT and Wnt signaling, both of which exert pro-metastatic effects that impair treatment responses^117^. Of particular interest were enriched targets of two oncogenic fusion proteins, *SS18-SSX* (misattributed in the GTRD database as *IGLV5-37* as noted above) and *SET-NUP214*, whose regulatory targets were enriched or differentially activated in association with both ColX module expression and signatures for osteogenesis and pathological OA cell types, suggesting common transcriptional dysregulation between these disease states.

Together, these results demonstrate that the genes in these cancer-specific ColX modules are involved in a diverse array of biological functions which may exacerbate the pathophysiology of both cancer and OA. Numerous genes implicated in both OA pathology and metastasis co-modularize with *COL10A1* in tumors, including *COMP*, *CXCL12*, and *FN1*, further suggesting common EMT-mediated dysregulation across these disease states. ColX module expression thus appears to highlight breast and pancreatic tumor states marked by activation of developmental and pro-OA pathways which may contribute to heightened metastatic potential and poorer clinical outcomes.

Survival analysis of the two TCGA cohorts reveals the prognostic value of ColX and its associated modules in both breast and pancreatic cancer. Increased expression of stromal ColX has previously been shown to correlate with worse survival outcomes in breast cancer cohorts with diverse tumor mutational burdens^13,118^, and *COL10A1* has also been linked to negative prognosis in pancreatic cancer^93^. Here, we corroborated that increased ColX expression confers significantly increased overall survival risk in both breast and pancreatic cancer. High ColX module expression is also significantly associated with decreased DFI in pancreatic cancer, an effect which is stronger in the male PAAD cohort compared to the female PAAD cohort. We observed that DFI events actually occurred more frequently in the high-ColX module group in the female PAAD cohort (65% event rate) compared to the male PAAD cohort (27% event rate), but the groupwise Kaplan-Meier analysis for the female PAAD cohort was ultimately not significant, likely due to being underpowered (when the analysis was rerun with all female samples artificially duplicated, the difference between high- and low-ColX module expression groups became significant at an α-level of 0.05). Analysis of a larger sample cohort would help clarify whether this is truly a gender-specific effect or merely the result of the cohort size analyzed here. Of note, a prior meta-analysis of multiple studies comprising 1,000 pancreatic cancer patients in total found that increased EMT is vital to the process of tumor budding, which confers significantly worse outcomes in terms of both overall survival and disease-free survival^109^. Thus, our results corroborate earlier findings regarding the predictive value of ColX expression and suggest the clinical utility of the ColX modules for risk stratification and prognostication.

Although *COL10A1* has been reported to be expressed by pancreatic cancer cells directly^73^, we observed that ColX is most strongly detected in the stromal region of PAAD tumors (Figure 1B), suggesting that it more likely serves as a marker for infiltration of fibroblasts and subsequent induction of an immunosuppressive microenvironment. Additionally, stromal markers such as *ACTA2* are co-modularized with *COL10A1* in all 4 ColX modules. These observations are consistent with *COL10A1*’s previously-characterized role as a marker of myofibroblastic TGF-β-driven fibroblasts in pancreatic cancer, based on single-cell analysis of human PDAC samples^78,119^. Recent work by Thorlacius-Ussing et al. has further highlighted *COL10A1* (as well as *COL8A1*, *COL11A1*, and *COL12A1*, all of which are present in the ColX modules) as a marker of myCAFs in PDAC as well as in other cancer types^93^. In tumors, the ECM is primarily produced by fibroblasts and activated myofibroblasts, the latter of which are additionally responsible for driving fibrosis in response to inflammatory stimuli such as TGF-β signaling from immune cells^1^. While CAFs comprise a heterogeneous collection of cells with a diverse array of functions, as a whole they exhibit numerous shared features across breast, pancreatic and other cancers, notably plasticity of their developmental pathways as well as purported regulatory roles in the functioning of NK, T, and other immune cells^120^. In breast and pancreatic tumors with high ColX module expression, we observed significant differential activation of numerous CAF types including matrix CAFs, inflammatory CAFs, antigen-presenting CAFs, vascular CAFs, and tumor-like CAFs, coupled with a decreased signal associated with dividing CAFs especially in breast cancer. This activation of CAF-specific gene signatures was corroborated by co-modularization of key CAF type markers with *COL10A1* across the breast and pancreatic ColX modules, which highlight CAF heterogeneity as well as common features of CAF-mediated aggression in both cancer types. CAFs have been shown to exert a variety of tumor-protective effects through direct interactions with cancer cells, ranging from promoting cancer stemness and preventing T cell recognition of cancer cells to facilitating invasion of the basement membrane through deposition of collagen “migratory tracks” and integrin-mediated interactions with fibronectin^1^. Although CAF populations are often characterized by their dominant role, considerable diversity of function has also been reported for several specific subtypes^19^. For example, although matrix CAFs are primarily responsible for producing and remodeling the ECM, they also appear to be capable of producing pro-inflammatory cytokines and chemokines to facilitate adhesion and migration. Similarly, tumor-like CAFs typically mimic tumor expression patterns and interact directly with tumor cells to promote stemness, chemoresistance and immunosuppression, but may also produce MMPs and other matrix proteins which contribute to remodeling. Our observed activation of antigen-presenting CAFs in pancreatic cancer is of particular interest, as prior studies in pancreatic cancer have shown that this CAF subtype interacts significantly with tumor-infiltrating T cells and TAMs through MHC II expression which induces naive CD4^+^ T cells to differentiate into regulatory T cells, thereby contributing to immune evasion^120,121^. Regulatory T cells and TAMs were among the most strongly activated immune cell types in high-ColX module pancreatic (as well as breast) tumors in our analysis, an effect which links high ColX module expression to both CAF activity and immunosuppression in aggressive tumors. Of note, the activation of regulatory T cells in the absence of activated CD8^+^ T cell signatures is an effect which is almost uniquely associated with the ColX modules (as opposed to other WGCNA modules), suggesting predominance of a pathological immunosuppressive environment which lacks a strong cell-mediated immune component. Additionally, recent work has shown that increased CAF density is associated with an inflammatory, pro-EMT environment in PDAC^122^. *COL10A1* is thus a valuable biomarker for CAF-mediated ECM remodeling, immunosuppression, inflammation, and induction of invasion in breast and pancreatic tumors; future work will further elucidate its complex roles across diverse CAF and cancer types.

Activated CAFs have been shown to derive from various stromal origins including bone marrow^1^. The upregulation of the TGF-β-induced *TGFBI* in BMSCs (Table S6) supports the hypothesis that ColX may originate from “activated,” dysregulated bone marrow-derived mesenchymal cells which infiltrate breast and pancreatic tumors. There is evidence for this effect in mouse models^123^, but effective markers are still being sought for human tumors. ColX is the strongest potential marker so far due to its specific expression in bone marrow during adulthood. We found that increased ColX module expression correlated significantly with the BMSC signature, as evidenced by CIBERSORTx analysis and co-modularization of *COL10A1* with BMSC-associated collagens type I, IV, VIII, XI, and XII. The roles that such bone-derived cells play within the tumor microenvironment is an area of active study. BMSCs are intricately involved in regulation of osteoblastic differentiation and the vascular microenvironment, and OA-MSCs, which are also ColX-positive, drive the chronic fibrosis and inflammation which characterizes the disease state^10,124^. In the context of solid tumors, high *COL10A1* expression may therefore indicate the presence of bone marrow-derived cells which contribute to an inflammatory, pro-EMT microenvironment. Coupled with the enrichment for gene ontologies related to cartilage/skeletal development and ossification, this suggests that increased ColX module expression may signify development of ECM pathology within the tumor mesenchyme. A compelling piece of evidence for this common developmental feature is the fact that the Wnt family gene *WNT2* emerged as a ColX module gene in both BRCA and PAAD (Table S2A) as well as a specific marker of BMSCs (Table S6); the Wnt signaling cascade is heavily implicated in joint degeneration and OA pathogenesis^125^, and *WNT2* has been established as a highly-expressed gene in breast cancer as well as a pro-metastatic activator in pancreatic cancer^126,127^. The interplay between various CAF subtypes and bone marrow cells is further supported by the co-modularization of *COL10A1* with key markers such as *CXCL12*, which is a marker for both inflammatory CAFs as well as hypertrophic chondrocytes which give rise to osteoblasts and recruit vasculature during skeletal development^19,128^. Corroborated by the capacity of NCSCs to transform into diverse pathological OA cell types through a “senescence-associated cell transition and interaction” (SACTAI)^124^ which mimics EMT-associated changes occurring in breast and pancreatic cancer, these results further support the idea that advanced tumors may contain significant subpopulations of cells with OA character. Additionally, the mutual enrichment of specific genes implicated in EMT across breast and pancreatic cancer and pathological OA cell types suggests that similar dysregulatory mechanisms are crucial to development of both disease states. Spatial transcriptomic studies would provide greater insight to the specific tumoral regions exhibiting *COL10A1* expression and help to further validate this hypothesis. Thus, differential activity of the ColX module reveals a more complete picture of the contribution of *COL10A1* to malignant CAF and pro-metastatic activity in advanced tumors.

Although our findings suggest a multifaceted role for *COL10A1* in the maintenance and remodeling of the ECM with involvement of CAFs, immune cells, and other key players in the tumor microenvironment, they are necessarily limited by the descriptive nature of this study. In particular, while we have highlighted numerous pathological mechanisms which appear to correlate and trend directionally with ColX module expression, direct experimentation will be necessary to robustly validate these relationships in an in vitro or in vivo setting. Survival analyses (in particular for the gender-specific PAAD cohorts) were restricted by the number of TCGA samples with complete data for each outcome of interest, and the ColX module-based survival impacts presented here would benefit from validation in a larger dataset. Nevertheless, the inflammatory, immunosuppressive, and pro-metastatic pathways we have identified here supplement the current knowledge regarding *COL10A1* and its significance to solid tumors, and suggest multiple avenues of exploration to further characterize its importance in breast, pancreatic, and other cancers.

## CONCLUSIONS

In this study, we have demonstrated numerous links of biological and clinical interest between the stromal ECM in breast and pancreatic cancer as well as bone marrow and OA cartilage, highlighted by shared expression of *COL10A1* and its associated gene networks which contribute to development of an inflammatory, immunosuppressive, and CAF-dominated microenvironment to facilitate EMT and metastasis. *COL10A1* is an important and specifically expressed collagen whose role in the progression of solid tumors is an area of active study; our results suggest that it holds substantial value as a regulator and biomarker of aggressive tumor phenotypes with implications for ECM-targeted therapies and clinical outcomes. Identification of tumors which exhibit high expression of *COL10A1* and its associated genes may reveal the presence of more aggressive pathological microenvironments with heightened EMT and metastatic potential. These findings may enable more effective risk assessment and treatment of patients with breast and pancreatic cancer.

## LIST OF ABBREVIATIONS

BH: Benjamini-Hochberg

BMSCs: bone marrow stromal cells

BRCA: breast invasive carcinoma

CAFs: cancer-associated fibroblasts

ColX: collagen type X (*COL10A1*)

CMT: chondrocyte-mesenchymal transition

DFI: disease-free interval

ECM: extracellular matrix

EMT: epithelial-to-mesenchymal transition

%G.A.M.E.: percentage of Genes Above-Median Expression

GEO: Gene Expression Omnibus

GO: gene ontology

GSEA: Gene Set Enrichment Analysis

GTRD: Gene Transcription Regulation Database

IHC: immunohistochemistry

ME: module eigengene

NCSCs: normal cartilage stromal cells

OA: osteoarthritis, osteoarthritic

OA-MSCs: OA mesenchymal stromal cells

OACs: OA chondrocytes

OS: overall survival

PAAD: pancreatic adenocarcinoma

PACA-AU: Pancreatic Cancer Australian

PDAC: pancreatic ductal adenocarcinoma

QuSAGE: Quantitative Set Analysis for Gene Expression

SACTAI: senescence-associated cell transition and interaction

TAMs: tumor-associated macrophages

TCGA: The Cancer Genome Atlas

TF: transcription factor

TFTs: transcription factor targets

WGCNA: weighted gene co-expression network analysis

## DECLARATIONS

### Ethics approval and consent to participate

Use of patient material was approved by the Lifespan institutional review board approval (IRB #1070389–9). All procedures were performed in accordance with the relevant guidelines and regulations.

### Consent for publication

Patient consent for sample publication was obtained appropriately.

### Availability of data and materials

The datasets supporting the conclusions of this article are available in the NCI Genomic Data Commons (https://gdc.cancer.gov/about-data/publications/pancanatlas), in the Broad Institute GDAC FireBrowse portal (http://firebrowse.org/), and in the GEO accession GSE176199 (https://www.ncbi.nlm.nih.gov/geo/query/acc.cgi?acc=GSE176199).

## Competing interests

ASB and MBR are co-inventors of the following patent: Brodsky, Alexander S. and Wang, Yihong and Resnick, Murray. 2017. Collagens as markers for breast cancer treatment. US Patent US09784743B2, filed Jun 20, 2016, and issued Oct 10, 2017. The other authors declare that they have no competing interests.

## Funding

This study was supported by R01AG080141 and P30GM122732.

## Authors’ contributions

EHHFY and AFY contributed equally. ASB and QC conceived and designed the study. WL generated the cartilage RNA-Seq data. DY performed IHC. EYW and AS interpreted IHC. MBR contributed to interpretation of all *COL10A1* and tumor features. EHHFY and AFY acquired, analyzed and interpreted the data, and drafted the manuscript. ASB and QC edited the manuscript with input from all authors.

## Acknowledgements

The results described here are in part based upon data generated by the TCGA Research Network (https://www.cancer.gov/tcga). We thank the patients and their families for their participation in the individual TCGA projects.

## SUPPLEMENTAL FIGURES

**Figure S1:** TCGA ColX modules are preserved in cancer microarray datasets. **(A and B)** Module preservation scores (*Z_summary_*) for all TCGA RNA-Seq-derived **(A)** breast and **(B)** pancreatic cancer WGCNA modules in comparably-sized microarray tumor datasets. ColX modules are indicated by **(A)** pink or **(B)** orange bars. Dotted lines indicate “high preservation” threshold of *Z_summary_* = 10 as defined by the authors of WGCNA.

**Figure S2:** TCGA WGCNA modules are enriched for numerous hallmark pathways. **(A–D)** Bubble plots of MSigDB hallmark pathway gene sets^54^ enrichment in WGCNA modules from **(A)** breast cancer, **(B)** pancreatic cancer, **(C)** male pancreatic cancer, and **(D)** female pancreatic cancer cohorts. ColX modules for each dataset are indicated by bolded labels. See Table S4 for significance values and Figure 2D for ColX module-specific enrichment results. **(E and F)** Overlap of EMT hallmark pathway genes within ColX WGCNA modules from TCGA **(E)** breast and pancreatic cancer and **(F)** gender-segregated pancreatic cancer datasets. Genes comprising each sector are listed alphabetically.

**Figure S3:** The G.A.M.E. metric effectively proxies ColX module expression and improves sample stratification. **(A–D)** Relationship between ColX module eigengene (ME) expression and proportion of “Genes Above Median Expression” (G.A.M.E.) for **(A)** breast cancer, **(B)** pancreatic cancer, **(C)** male pancreatic cancer, and **(D)** female pancreatic cancer cohorts. Colored curves represent densities of ME (*right*) and G.A.M.E. (*top*) variables for each panel. Blue dotted lines indicate Jenks natural breakpoints defining 3 clusters (“low”, “medium”, and “high”). Spearman correlations (*ρ*) are shown for each panel.

**Figure S4:** TCGA WGCNA module expression tracks differential activity of immune, transcription factor, and cancer-associated fibroblast signatures. **(A–P)** Mean pathway activation (by QuSAGE) of various gene sets in samples with high module expression relative to samples with low module expression for each WGCNA module and TCGA cohort. **(A–D)** Results for cancer immunome genesets in **(A)** breast cancer, **(B)** pancreatic cancer, **(C)** male pancreatic cancer, and **(D)** female pancreatic cancer cohorts. Gene sets within each block are ordered by significance in respective ColX module; see Figure 3B for focused comparison. **(E–H)** Results for transcription factor target lists in **(E)** breast cancer, **(F)** pancreatic cancer, **(G)** male pancreatic cancer, and **(H)** female pancreatic cancer cohorts. Gene sets within each block are ordered by effect size in respective ColX module; see Figure 3C for focused comparison. **(I–P)** Results for cancer-associated fibroblast (CAF) gene sets derived from **(I–L)** breast tumor scRNA-Seq data and **(M–P)** pancreas/other tumor scRNA-Seq data, in **(I and M)** breast cancer, **(J and N)** pancreatic cancer, **(K and O)** male pancreatic cancer, and **(L and P)** female pancreatic cancer cohorts. Gene sets within each block are ordered by significance in respective ColX module; see Figures 3D–E for focused comparison. See Methods section for sample grouping process (note the same procedure for calculating G.A.M.E. and assigning “low” and “high” labels was applied to each WGCNA module respectively). Significance values were corrected across each module individually (by rows). Significance values: *, p.adj < 0.05; **, p.adj < 0.01; ***, p.adj < 0.001; ****, p.adj < 0.0001.

**Figure S5: TCGA WGCNA module expression correlates with variable survival risk. (A–D)** BH-adjusted significance values for multivariate Cox proportional hazards models conditioning overall survival (OS), disease-specific survival (DSS), disease-free interval (DFI), or progression-free interval (PFI) on age, gender, binarized tumor stage, and module eigengene (ME) expression for WGCNA modules from **(A)** breast cancer, **(B)** pancreatic cancer, **(C)** male pancreatic cancer, and **(D)** female pancreatic cancer cohorts. ColX modules for each dataset are indicated by bolded labels (see Figure 4B for focused ColX module results). Significance values: *, p.adj < 0.05; **, p.adj < 0.01; ***, p.adj < 0.001; ****, p.adj < 0.0001.

**Figure S6: Osteoarthritis cell type-specific markers are enriched for ColX module genes. (A–D)** Bubble plots of OA cell type-specific marker gene enrichment in WGCNA modules from **(A)** breast cancer, **(B)** pancreatic cancer, **(C)** male pancreatic cancer, and **(D)** female pancreatic cancer cohorts. ColX modules for each dataset are indicated by bolded labels. **(E)** Overlap of genes within breast and pancreatic cancer ColX modules and OA cell type-specific gene sets. **(F)** Overlap of EMT pathway genes within breast and pancreatic cancer ColX modules and OA cell type-specific gene sets. Unlabeled sectors represent 0 gene overlap. NCSC is not shown as no NCSC-specific markers overlap with EMT pathway genes. NCSC, normal cartilage stromal cells; OA-MSC, osteoarthritis mesenchymal stromal cells; OAC, osteoarthritis chondrocytes; BMSC, bone marrow stromal cells.

**Figure S7:** Collagen gene expression varies across bone and cartilage cell types and TCGA cohorts. **(A)** Normalized expression of all collagen genes expressed at nontrivial levels in bone marrow and cartilage cell types. Note that *COL6A4P1*, *COL6A5*, *COL20A1*, and *COL23A1* were filtered out as “low-expression” genes and are omitted here. NCSC, normal cartilage stromal cells; OA-MSC, osteoarthritis mesenchymal stromal cells; OAC, osteoarthritis chondrocytes; BMSC, bone marrow stromal cells. **(B)** Normalized expression of all collagen genes expressed at nontrivial levels in TCGA cohorts. Note that *COL6A4P1*, *COL6A5*, *COL20A1*, and *COL26A1* were filtered out as “low-expression” genes and are omitted here; additionally, *COL2A1* was only nontrivially expressed in BRCA.

## SUPPLEMENTAL TABLES

**Table S1:** Overview of cancer datasets used in this study. **(A)** Breast and pancreatic cancer datasets used in this study. **(B)** Citations of breast cancer GEO accessions used in this study.

**Table S2:** Gene modules characterized in this study. **(A)** WGCNA-generated *COL10A1* (ColX) modules from TCGA breast and pancreatic cancer datasets, as well as gender-specific pancreatic cancer datasets. **(B)** WGCNA module assignments for all expressed genes in each dataset. Modules are numbered in descending order of size, beginning with module 1. “-” indicates genes which were filtered out based on low expression; genes labeled with a 0 were not assigned to any module by WGCNA.

**Table S3: Gene ontology and Reactome pathway enrichment analysis of ColX modules. (A–E)** Gene ontology pathway enrichment for **(A)** breast cancer, **(B)** pancreatic cancer, **(C)** overlap between breast and pancreatic cancer, **(D)** male pancreatic cancer, and **(E)** female pancreatic cancer ColX modules. **(F–J)** Reactome pathway enrichment for **(F)** breast cancer, **(G)** pancreatic cancer, **(H)** overlap between breast and pancreatic cancer, **(I)** male pancreatic cancer, and **(J)** female pancreatic cancer ColX modules. All significantly enriched Reactome pathways/GO terms (q < 0.05) are shown.

**Table S4:** Hallmark pathway enrichment analysis of WGCNA modules. **(A–D)** Hallmark pathway enrichment for **(A)** breast cancer, **(B)** pancreatic cancer, **(C)** male pancreatic cancer, and **(D)** female pancreatic cancer ColX modules. All p-values shown were BH-corrected across all 50 hallmark pathways for each module. ColX modules for each dataset (#8, #13, #7, and #23, respectively) are shown in the first column for clarity. Related to Figure S2A–D.

**Table S5:** TFs implicated in ColX modules. **(A)** GTRD TFTs enriched in BRCA and PAAD ColX modules. Cancer-relevant TF functions and overlapping targets are shown where applicable. (Related to Figure 2E.) **(B)** Selected QuSAGE-significant transcription factors whose targets are differentially activated/inactivated between tumors with high and low ColX module expression. Cancer/OA-relevant TF functions are shown where applicable. (Related to Figure 3C.)

**Table S6: Differential expression of bone marrow and cartilage cell type-specific genes.** List of all genes defined as specific to normal cartilage stromal cells/NCSCs, OA mesenchymal stromal cells/OA-MSCs, OA chondrocytes/OACs, and bone marrow stromal cells/BMSCs, based on OA RNA-Seq data. Differential expression results were computed across all cells and statistics were extracted for each pairwise comparison. See Methods section for definition of “cell type-specific” genes.

## Notes

https://gdc.cancer.gov/about-data/publications/pancanatlas

http://firebrowse.org/

https://www.ncbi.nlm.nih.gov/geo/query/acc.cgi?acc=GSE176199

